# Learning gene co-regulation across cells with Multicellular Transformer

**DOI:** 10.64898/2026.11.14.766251

**Authors:** Maciej Wiatrak, Alma Andersson, Kam Hon Hoi, Tommaso Biancalani, Aïcha BenTaieb

## Abstract

Understanding how genes coordinate across cells is central to explaining tissue function and disease, yet most existing models still take a single cell as input and therefore emphasize cell-intrinsic regulation. We present the Multicellular Transformer (MCT), a foundation model built around a different assumption: cells function together within tissues, so the appropriate unit of input is not one cell in isolation but a set of cells from the same sample. By jointly modeling multiple cells, MCT learns context-aware gene embeddings that capture gene-gene interactions across the tissue environment. Trained on a large set of tissue samples (>4200) across more than 30 tissue types, MCT recovers ligand-receptor interactions, infers cross-cell signaling programs, and reveals spatial co-localization patterns directly from scRNA-seq data. Its contextual embeddings identify disease-specific rewiring of gene neighborhoods in systemic lupus erythematosus and COVID-19 and predict population-level perturbation effects in cancer without additional training. By linking gene-level regulation to tissue-scale organization, MCT provides a scalable framework for modeling multicellular biology and enables *in silico* discovery of therapeutic targets and tissue-specific dependencies.

## 1. Introduction

The success of foundation models in language and vision has spurred parallel efforts in single-cell biology. However, most existing models such as GeneFormer, scGPT and scPRINT adopt a cell-centric perspective (Theodoris et al. 2023; Cui et al. 2024; Kalfon et al. 2025), taking a single cell as input to learn representations that generalize across downstream tasks. While recent methods (Wen et al. 2023; Schaar et al. 2024) have begun to incorporate tissue context, they remain rooted in the single-cell paradigm. Consequently, their underlying gene embeddings do not fully reflect the dynamic shifts in gene function that occur across diverse cellular or tissue environments.

We address this limitation with the Multicellular Transformer (MCT), a foundation model built on the principle that the sample, rather than the isolated cell, is the fundamental unit of tissue function. MCT treats the full population of cells within a tissue as a single observation, representing samples as collections of contextualized gene embeddings learned jointly. This design acknowledges that a cell’s state is defined not only by its intrinsic identity but by its surrounding multicellular environment; for instance, the regulatory program of a T cell is fundamentally shaped by whether it resides in a tumor-rich niche or a lymph node.

Current computational approaches to modeling gene coordination remain limited in scaling and generalization. Frameworks like SCENIC (Sande et al. 2020) and CellOracle (Kamimoto et al. 2023) prioritize cell-intrinsic regulatory networks, while cross-cell methods like DIALOGUE (Jerby-Arnon and Regev 2022) often rely on correlation-based modules that may not generalize across heterogeneous tissues. Methods that process cells independently are inherently poorly matched to phenomena like coordinated inflammation or tissue remodeling—which only become visible when cells are modeled as a collective system.

MCT fills this gap by utilizing an architecture and training strategy specifically designed to capture cross-cell regulatory interactions. By training on a corpus of 14 million cells from over 3,500 donors across 4,200 samples, MCT learns representations where co-regulated genes cluster in a context-aware latent space. We demonstrate that this approach enables unsupervised receptor-ligand prediction, the recovery of spatially-informed interaction patterns, and zero-shot population-level perturbation analysis. By shifting from single-cell to sample-level input, MCT provides a framework for biomarker discovery and therapeutic target prioritization that reflects the actual complexity of multicellular biology.

## 2. Results

### 2.1. Learning gene co-regulation across cells with Multicellular Transformer

Cells within a tissue do not act independently: they coordinate through context-dependent signaling mediated by ligands, receptors, and other communication factors. To model biology at the level at which tissues actually function, we developed the Multicellular Transformer (MCT). In contrast to existing single-cell foundation models such as GeneFormer, scGPT and scPRINT (Theodoris et al. 2023; Cui et al. 2024; Kalfon et al. 2025) which model only a single-cell, MCT takes multiple cells from the same sample as input and learns gene representations from their joint expression context. Specifically, MCT takes as input a set of single-cell profiles – from scRNA-seq or spatial transcriptomics data – derived from the same tissue sample. These profiles form a cell-by-gene matrix *X* ∈ **R***^M^*^×^*^N^*, where *M* and *N* denote the number of cells and genes. Instead of encoding each cell separately, MCT treats the sample as a coordinated multicellular system. Each gene is represented by a latent embedding that combines a sample-invariant gene function embedding with an expression embedding derived from that gene’s expression across all input cells. The transformer then contextualizes these representations by allowing genes to exchange information with one another, so each gene embedding reflects not only its own expression but also the surrounding cellular composition of the sample.

The gene expression latent vector is computed with a shared two-layer feed-forward neural network (FFNN), which reduces overfitting. During pretraining, its input units are fixed to specific cell types according to their abundance in the dataset (Methods); for example, units 1–20 are assigned to *CD8 T cells*, whereas units 21–32 correspond to *Macrophages*. This design makes the encoder cell-type aware and discourages it from learning only average expression patterns. We examine sensitivity to cell resampling and within-cell-type ordering in Appendix C, and demonstrate that contextual embeddings are robust to both. The gene function embedding projects gene identity into a context-independent vector. Both components are learned from scratch and summed to form the final gene embedding, so that co-regulated genes become close in latent space.

For pretraining, we compiled a corpus of over 14 million cells from more than 3,500 donors spanning diverse tissues and diseases, harmonizing cell types across datasets (Supp. Fig. S1). MCT uses a 12-layer encoder and nearly 60,000 genes. To mitigate overfitting across sample observations, we apply a dynamic data augmentation strategy that iteratively sub-samples cells from each sample during training, see Appendix A.2. Because it considers all genes expressed in a sample rather than only those expressed in one cell, its context is much larger than that of single-cell foundation models such as scGPT (Cui et al. 2024) and GeneFormer (Theodoris et al. 2023). Importantly, the model can work with any sample where a subset of the genes seen during pre-training are present, making it highly flexible and adaptable to new samples. This larger context improves downstream performance (Fig. S2b). MCT learns gene co-regulation across cells by predicting masked gene identities from expression and context (Fig. 1d). We pair this masked objective with a reconstruction loss that predicts each gene’s expression from its contextual embedding, encouraging richer latent representations (Methods). We also provide insights on the MCT model’s scaling laws under varying data sizes in Appendix D.

**FIGURE 1.**
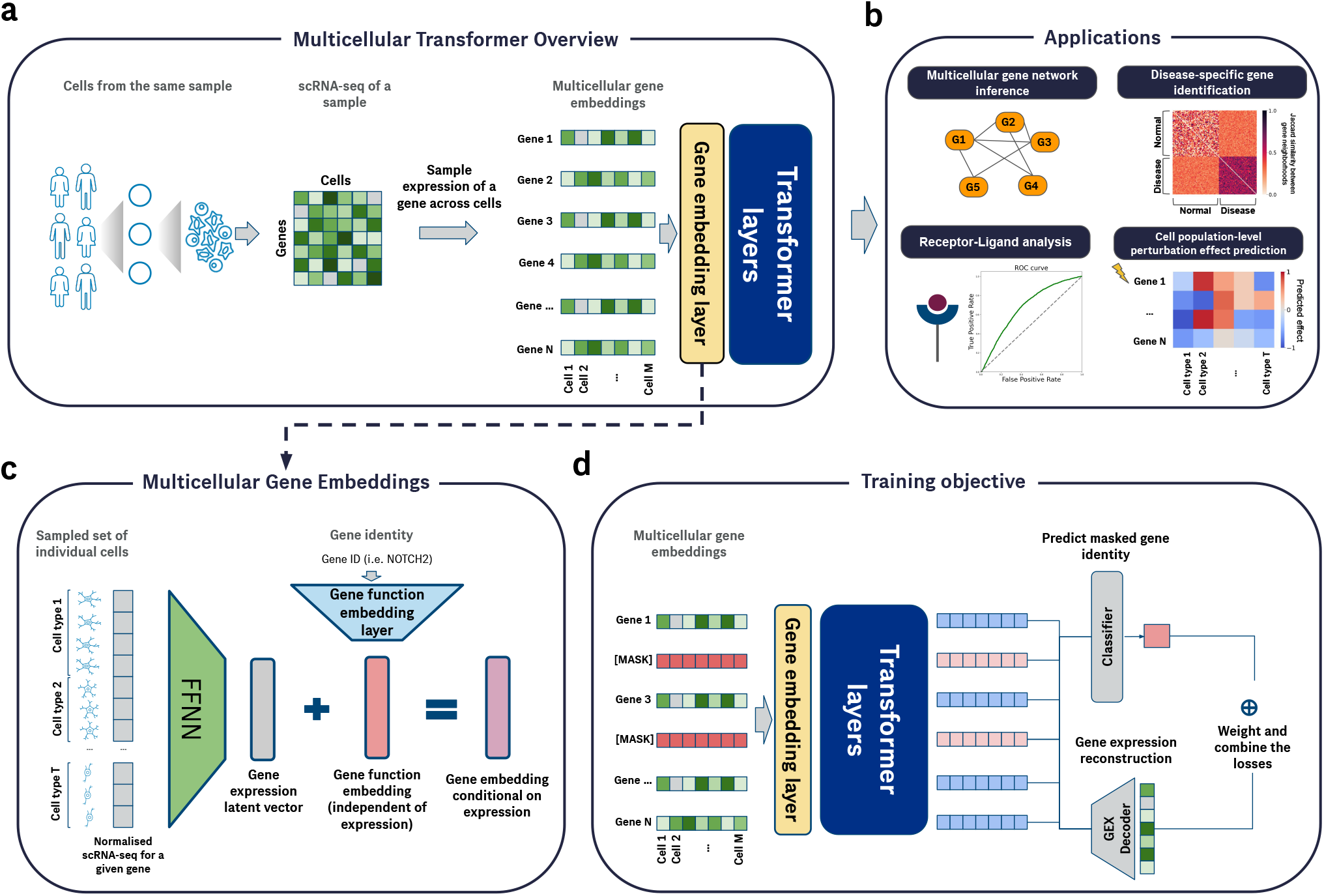
Multicellular Transformer learns gene co-regulation in a tissue environment. **(a)** Multicellular Transformer takes as input scRNA-seq from a set of cells originating from the same sample. The normalized expression of each gene across cells is used to compute gene embeddings, which are then passed to the transformer. The model produces contextual gene embeddings across cells in a sample that can be used in downstream tasks. **(b)** Multicellular Transformer can be applied to several downstream tasks, including gene network inference, disease-specific gene identification, receptor-ligand analysis, and zero-shot perturbation effect prediction at the cell-population level. **(c)** Multicellular gene embeddings. The expression of each gene across cells is fed into a feed-forward neural network (FFNN), which is shared across all genes to prevent overfitting. To capture interactions between cell types, the FFNN input units are fixed to specific cell types during pretraining. The final gene embedding consists of a gene expression latent vector, which varies with expression, and a gene function embedding, which is independent of expression. The gene expression latent vector captures the activity of the gene in the sample, whereas the gene function embedding represents its sample-independent function and is learned from scratch. The gene embedding layer and gene expression FFNN are summed to form the final embedding used as input to the transformer. **(d)** Multicellular Transformer is trained with a masked learning objective, randomly masking 15% of gene function embeddings and/or expression values, together with a gene expression reconstruction loss (Methods).

### 2.2. Multicellular Transformer uncovers gene interactions involved in cell-cell communication

#### Identifying receptor-ligand interactions

Ligand–receptor pairs provide a direct test of cross-cell gene relationships. In MCT, genes with similar embeddings are expected to participate in related interactions or processes, so we ranked candidate ligand–receptor pairs by cosine similarity between pretrained gene embeddings. To assess whether multicellular modeling improves this task, we benchmarked MCT against several cell-centric single-cell foundation models, including scGPT, scPRINT, CellFM, scFoundation and CellPLM (Cui et al. 2024; Kalfon et al. 2025; Zeng et al. 2025; Hao et al. 2024; Wen et al. 2023). Notably, this differs from traditional cell–cell communication methods, which rely on pre-defined databases to score known ligand–receptor pairs across cell populations; here, we use pretrained gene embeddings in a zero-shot setting to test whether any arbitrary pair of genes is likely to participate in a cross-cell process. Embedding similarity therefore reflects general interaction potential and does not, by itself, specify the underlying interaction type.

Against a consensus set of known receptor-ligand pairs from CellChatDB (Jin et al. 2021) (Methods), zero-shot gene embeddings from MCT outperformed cell-centric single-cell foundation models on AUROC and precision-recall metrics, reaching an AUROC of 0.71 versus 0.65 for the second-best method, scGPT (Cui et al. 2024) (Fig. 2a & Supp. Fig. S3). The gain held across distinct interaction types, including cell-cell contact, secreted signaling, and ECM-receptor interactions, as well as across distinct receptors (Supp. Fig. S4). To test whether better intercellular modeling came at the expense of intracellular relationships, we also evaluated transcription factor (TF)–target pairs from CollecTRI (Müller-Dott et al. 2023) and protein–protein interaction (PPI) pairs from OmniPath (Türei, Korcsmáros, and Saez-Rodriguez 2016). MCT performed competitively to other methods on these benchmarks (Supp. Fig. S5), indicating that it retains strong intracellular resolution.

**FIGURE 2.**
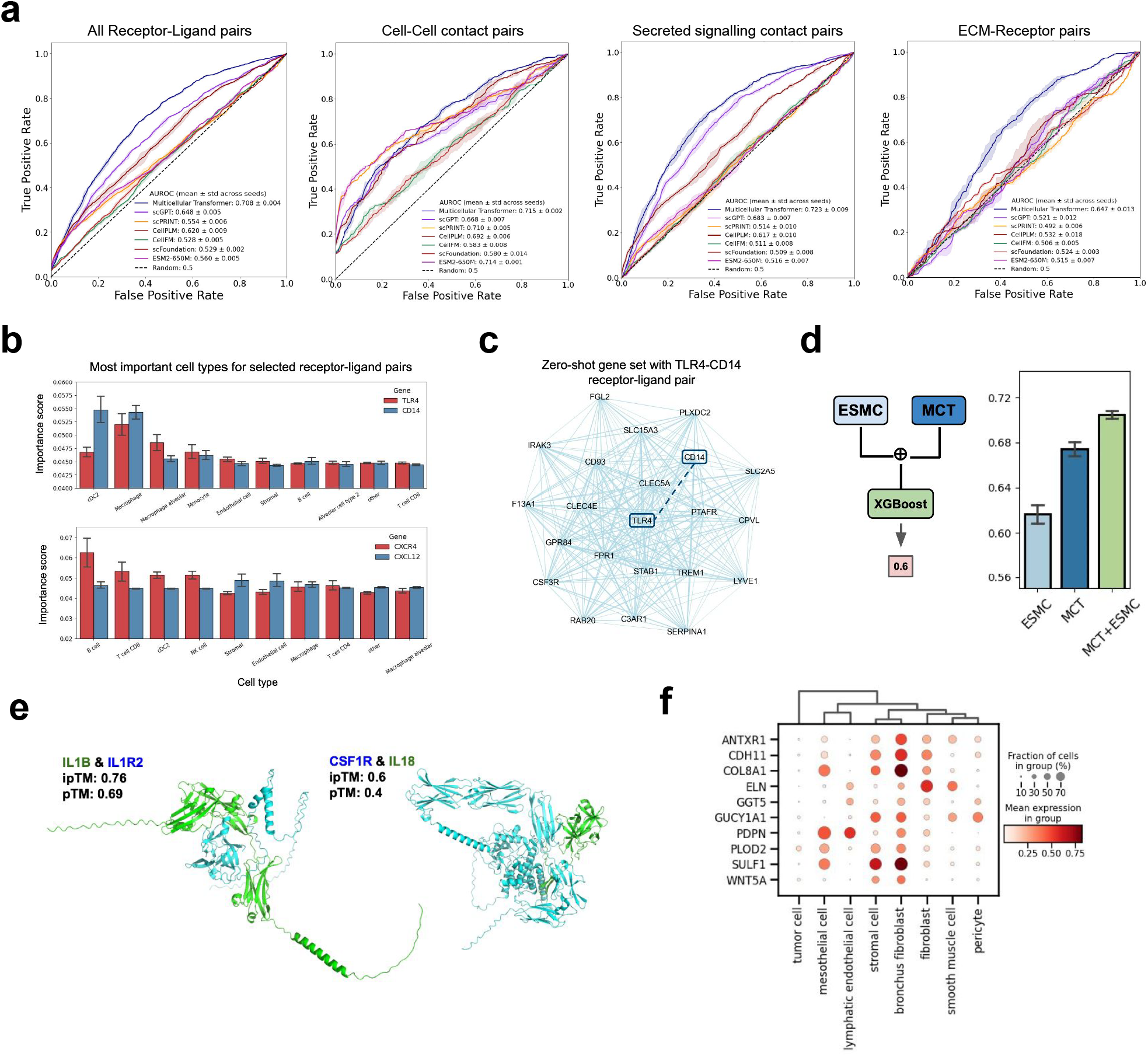
Multicellular Transformers identifies genes driving multicellular interactions. **(a)** Performance on the receptor-ligand prediction task using pretrained gene embeddings (zero-shot) across methods as measured by an area under the receiver operating characteristic curve (AUROC). The true receptor-ligand pairs were taken from CellChatDB receptor-ligand set Jin et al. (2021), while the negatives were randomly sampled with different seeds (Methods). Scores for each receptor-ligand pair were computed by calculating cosine similarity between embeddings coming from each model. We stratify the results the interaction type using the annotations from CellChatDB. **(b)** Most important cell types for *TLR*-*CD14* and *CXCR4*-*CXCL12* receptor-ligand pairs using integrated gradients (Sundararajan, Taly, and Yan 2017) on lung cancer atlas (Salcher et al. 2022). Cell type importance scores have been computed by applying softmax function to aggregated absolute attribution scores across samples. The error bars represent 95% confidence interval. cDC stands for conventional dendritic cells. **(c)** Example of a zero-shot gene set with a receptor (*CD14*) - ligand (*TLR4*) pair identified with Multicellular Transformer. **(d)** Combining transcriptomic and protein information from Multicellular Transformer (MCT) and ESM-C protein language model ESM Team (2024) leads to improved performance on receptor-ligand prediction task. Scores for each receptor-ligand pair were computed by calculating cosine similarity between embeddings coming from each model. The scores were then used as input to the XGBoost model which was trained and evaluated using 5-fold cross validation on the Liana consensus set (Methods). **(e)** Structures of protein complexes predicted with AlphaFold3 (Abramson et al. 2024) and predicted to interact with Multicellular Transformer. The pTM score measure how well different parts of protein fold together, while ipTM assesses the confidence of protein-protein interfaces, higher is better. **(f)** Normalized expression of a selected gene set involved in *Wnt* signaling in lung cancer across a set of cell types. For brevity we have limited cell types to the ones highly expressed in the gene set.

We hypothesized that this performance gain stems from modeling the full multicellular environment rather than only the two genes of interest. Consistent with this idea, AUROC increased with context length, underscoring the importance of multicellular context for receptor-ligand identification (Supp. Fig. S2).

To identify which cell types drive these predictions, we used integrated gradients (Sundararajan, Taly, and Yan 2017) (Fig. 2b). For the *TLR4*-*CD14* pair, MCT assigned the highest attribution to dendritic cells (cDC2) and monocytes, consistent with their role in antibacterial immune responses. For *CXCR4*-*CXCL12*, the model highlighted B and T cells together with stromal and endothelial cells, matching known roles in inflammation and angiogenesis (Liekens, Schols, and Hatse 2010). MCT also prioritized regulatory T cells for *IL2RA*-*IL2R* (Cheng, Yu, and Malek 2011) and tumor cells for *MET*-*HGF* (Cecchi, Rabe, and Bottaro 2010) (Supp. Fig. S6c).

Because signaling often depends on additional genes beyond the receptor and ligand themselves, we next examined gene sets extracted from pretrained MCT embeddings. These sets recovered known receptor-ligand-containing programs, including *TLR4*-*CD14* (Fig. 2c) and *WNT5A*-*FZD1*, part of the Wnt signaling pathway (Ring et al. 2014) (Supp. Fig. S6a).

#### Integrating multicellular and protein-level information

MCT predicts receptor-ligand interactions from scRNA-seq data alone, although the interaction itself is mediated by protein binding. To incorporate protein-level information, we combined MCT embeddings with those from the protein language model ESM-C (ESM Team 2024) in a boosting framework (Fig. 2d), which improved precision.

Because many human cell-surface receptors remain orphan and protein-binding simulations are expensive, we asked whether MCT could prioritize likely protein complexes. Using cytokine-focused lupus data (Perez et al. 2022), we ranked gene pairs by contextual embedding similarity, filtered for secreted–membrane pairs, and evaluated the top 70 candidates with AlphaFold3 (Abramson et al. 2024). Six pairs showed ipTM > 0.5 (mean = 0.26 ± 0.13), including known interactions such as *IL1B*-*IL1R2* (ipTM: 0.76) (Colotta et al. 1993) and *CSF1R*-*IL18* (ipTM: 0.6) (Ma et al. 2023) (Fig. 2e; Supp. Fig. S7a). These results suggest that MCT can help prioritize candidate receptor-ligand pairs for follow-up, although novel interactions still require experimental validation.

Having established that MCT recovers receptor-ligand relationships, we next examined broader co-regulatory programs learned during pretraining. We clustered pretrained gene embeddings to identify groups of co-regulated genes with similar latent representations (Methods).

Several gene sets overlapped with key pathways involved in multicellular signaling, including type I interferon, chemokine-mediated signaling, and T-cell receptor signaling (Supp. Fig. S8). In the lung cancer atlas (Salcher et al. 2022), these sets were expressed across diverse cell types (Supp. Fig. S9d), supporting the idea that MCT captures relationships spanning distinct cell states.

#### Revealing interactions promoting tumor progression in lung cancer

We then examined an MCT-derived gene set from the lung cancer atlas (Salcher et al. 2022). This set was enriched across tumor cells, fibroblasts, lymphatic endothelial cells, and other stromal compartments (Fig. 2f), and contained multiple genes linked to epithelial-mesenchymal transition (EMT), extracellular matrix remodeling, motility, and angiogenesis, including *CDH11*, *PDPN*, *WNT5A*, *PLOD2*, *COL8A1*, *ELN*, *SULF1*, and *ANTXR1* (Kiener et al. 2006; Wicki and Christofori 2007; Dissanayake et al. 2007; Hong et al. 2024; Lai et al. 2008; Carson-Walter et al. 2001). Their coordinated expression across mesothelial, fibroblast, pericyte, and endothelial compartments points to a multicellular program that supports tumor progression and metastasis.

### 2.3. Multicellular Transformer discovers disease-specific genes with context-aware gene embeddings

MCT produces sample-specific contextual embeddings, enabling the comparison of gene-gene interactions across states, such as *healthy* and *disease*. Since the neighbors of a gene’s embedding represent likely interaction partners, we used Jaccard similarity to compare top *k* neighborhoods across samples, identifying differential interactions between groups (Methods; Supp. Fig. S10). Genes with low overlap between conditions but high consistency within a condition are candidates for disease-specific rewiring. Interferon-stimulated genes such as *IFI44* and *OAS1*, for example, cluster with interferon-pathway genes in lupus and severe COVID-19 but not in healthy samples. This analysis consistently highlighted disease- and tissue-specific genes across SLE (Perez et al. 2022), COVID-19 (Stephenson et al. 2021), and multi-tissue data (Suo et al. 2022) (Fig. 3a; Supp. Fig. S11).

**FIGURE 3.**
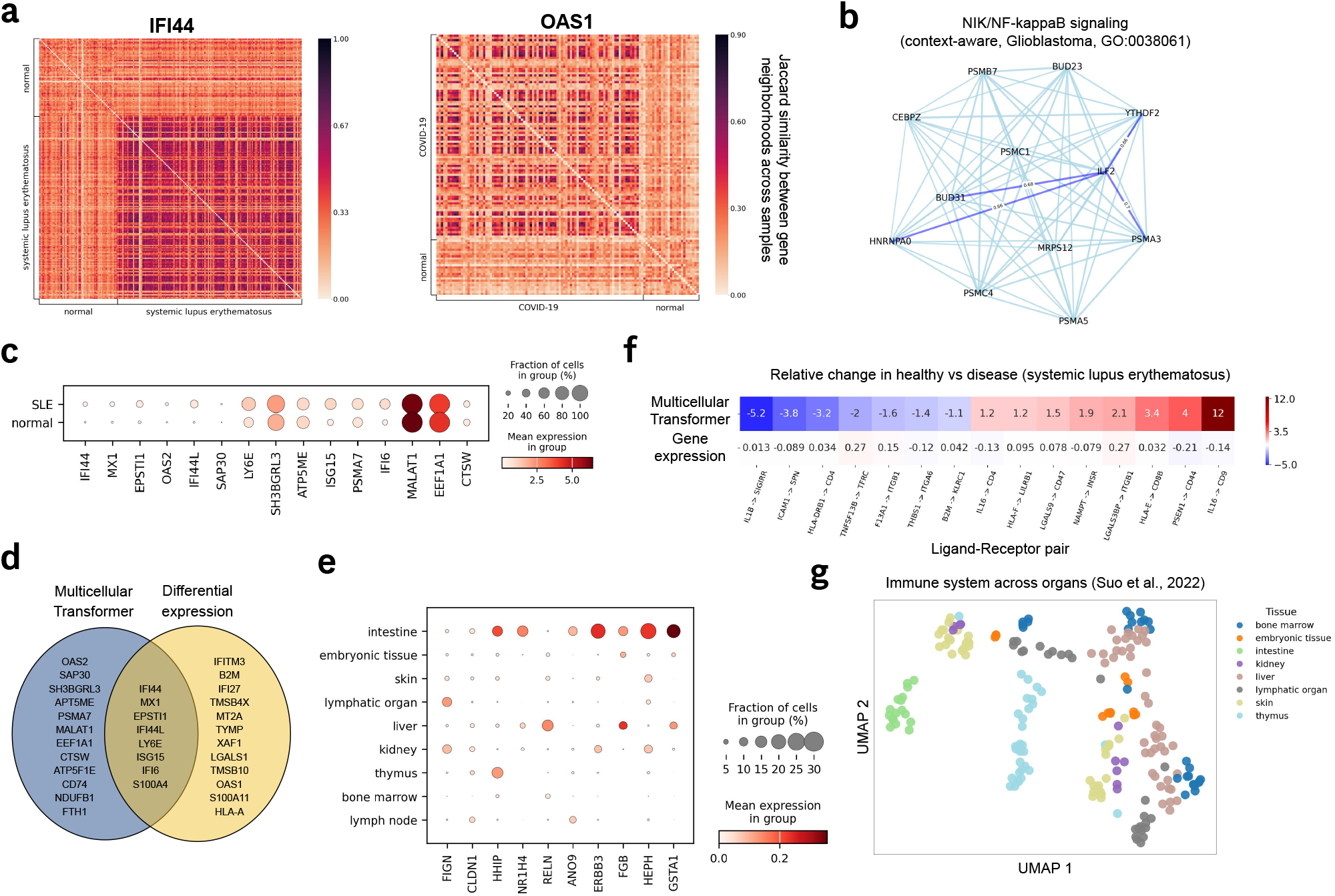
Context-aware gene embeddings from Multicellular Transformer identify disease- and tissue-specific genes. **(a)** Genes with high condition-specificity across SLE (Perez et al. 2022) & COVID-19 (Stephenson et al. 2021). The condition-specificity was calculated by dividing the mean overlap across distinct samples between gene neighborhoods in the same condition by the mean overlap between gene neighborhoods across conditions. **(b)** Example of a context-aware gene set enriched in *NF-*κ*B* pathway identified at Leiden resolution of 50 with Multicellular Transformer in a glioblastoma sample (Ruiz-Moreno et al. 2022). **(c)** The expression of disease-specific genes in SLE (Perez et al. 2022) computed using context-aware embeddings from Multicellular Transformer. Many genes have been shown to be associated with the disease but do not exhibit a significant change in expression between healthy and disease. **(d)** The overlap between top 20 genes discovered by the Multicellular Transformer and differential expression (Wilcoxon). Overlap 40%, *N* 20. **(e)** The expression of highly tissue-specific genes according to the gene neighborhood across samples computed with MCT contextual gene embeddings on the immune across organs dataset (Suo et al. 2022) across different tissues. **(f)** Relative change in cosine similarity (top) vs expression (bottom) in healthy vs disease patients with SLE across a set of known receptor-ligand pairs with highest change according to cosine similarity between MCT contextual gene embeddings. Receptor-ligand pairs with highest change according to expression can be found in Supp. Fig. S13b. **(g)** UMAPs of sample-level embeddings across tissues on the immune across organs dataset (Suo et al. 2022). The sample embeddings were extracted by averaging out all of the contextual gene embeddings in the sample using MCT.

We also clustered sample-specific embeddings from MCT’s last hidden layer to form context-aware gene sets. Glioblastoma-derived clusters were enriched for *NF-*κ*B*, *p53*, and stress-response pathways, whereas healthy peripheral blood mononuclear cell (PBMC) samples were enriched for canonical immune programs such as B-cell differentiation and T-cell receptor signaling (Fig. 3b; Supp. Fig. S12).

#### Identifying disease-specific genes in SLE and COVID-19

Applying this framework to systemic lupus erythematosus (SLE) and COVID-19 highlights genes whose roles shift despite modest expression changes. In lupus (Perez et al. 2022), *IFI44* and *MX1* (Fig. 3c) emerged as robust SLE-specific genes, consistent with their known links to interferon signaling (Rodríguez-Carrio et al. 2019; AlFadhli et al. 2016), whereas *OAS1* and *CD74* were identified as COVID-19-specific genes (Stephenson et al. 2021; Banday et al. 2022; Westmeier et al. 2023) (Supp. Fig. S13a).

Comparing MCT’s top 20 SLE-specific genes with those from differential expression shows a 40% overlap, confirming recovery of established biomarkers (Fig. 3d). At the same time, MCT identifies additional candidates such as *OAS2*, *FTH1*, and *SH3BGRL3*, whose neighborhoods shift even when expression changes are small. For example, *SH3BGRL3* associates with T-cell activation genes in lupus but not in healthy controls, and *OAS2* is flagged through a neighborhood enriched for IFN-inducible genes.

MCT’s context-awareness also extends to tissue specificity. Genes such as *FIGN*, *CLDN1*, and *GSTA1* from the immune system across organs dataset (Suo et al. 2022) remain clustered only in tissues where they are functionally relevant (Fig. 3e), showing that MCT captures subtle tissue-restricted roles without relying on large expression differences (Lavin et al. 2014).

#### Revealing receptor-ligand interactions in systemic lupus erythematosus

Many disease processes emerge from coordinated actions among multiple genes. To capture such changes, we used MCT to compare receptor-ligand similarity between healthy and disease states in SLE.

Traditional scRNA-seq analyses may miss lowly expressed or context-dependent interactions, yet MCT uncovered receptor-ligand pairs that shift strongly between healthy and SLE states even when expression remains stable (Fig. 3f & Supp. Fig. S13b). These included *IL16*-*CD9* and *IL1B*-*SIGIRR*, together with established lupus interactions such as *TNFSF13B*-*TNFRSF13C* (Lee et al. 1998; Jeffries et al. 2011; Song et al. 2014; Wang et al. 2013). MCT also grouped *CD9* with *OAS1*, *TNFSF10*, and *WARS1* (Supp. Fig. S13c), linking these shifts to immune regulation, inflammation, and apoptosis (Dinarello 2000; Postal and Appenzeller 2011; Gao, Tan, and Luo 2020; Kuribayashi et al. 2008; Scandura et al. 2004; Weedon 2007; Nguyen et al. 2020).

Finally, sample-level embeddings obtained by averaging gene embeddings clustered in a biologically meaningful way. Samples from different tissues or disease states separated clearly and outperformed expression-based baselines such as PCA as measured by normalized mutual information in 3 of 4 comparisons (Fig. 3g; Supp. Fig. S14), indicating that MCT yields interpretable sample-level structure as well as gene-level insights.

### 2.4. Multicellular Transformer allows for *in silico* zero-shot perturbation prediction at cell population level

Because MCT captures contextual gene interactions in multicellular environments, it should also be able to predict the effects of genetic perturbations on entire cell populations in a zero-shot setting. This is particularly relevant for assessing how therapeutic interventions affect both target cells and surrounding cell types. Most existing perturbation models focus on cell lines (Roohani, Huang, and Leskovec 2024; Lotfollahi et al. 2023; Cui et al. 2024) and therefore ignore the broader tissue microenvironment. In contrast, MCT enables *in silico* perturbations in multicellular samples by operating on contextual gene embeddings. Because population-level perturbation data are scarce, all analyses were performed without additional fine-tuning.

#### Uncovering multicellular gene dependencies with *in silico* genetic perturbations

We first tested whether MCT could recover cross-cell signaling programs and pathway dependencies under perturbation Using the fine-tuned model in the lung cancer atlas (Salcher et al. 2022), we simulated knockouts by removing a target gene from the input sample and comparing the predictions with the unperturbed baseline measured by the impact on the probability of predictions of gene identity (Methods).

MCT was shown to capture the dependencies between the *in silico* perturbed genes and their known interacting genes, achieving significantly higher knock-out effect compared to control genes (Fig. 4a; Supp. Fig. S15a; Methods), supporting its use for multicellular perturbation analysis.

**FIGURE 4.**
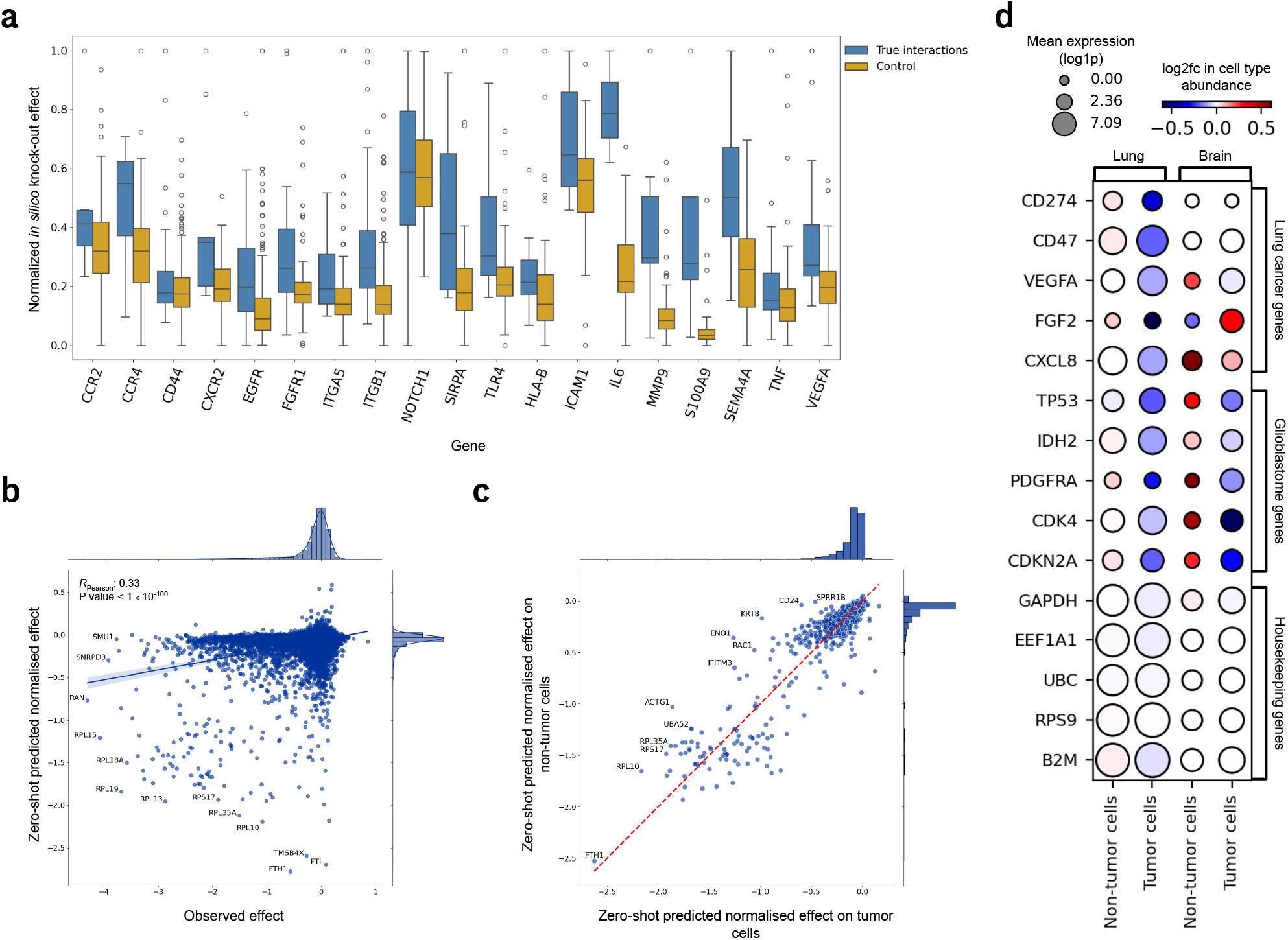
*In silico* genetic perturbation effect estimation at cell population level. **(a)** *In silico* gene knock-out effect on a set of true interacting genes and control genes. The control genes were randomly selected (Methods). The knock-out effect is quantified by the maximum log-fold change in gene’s logits (Methods). The predictions were min max normalized within a gene. The box represents the interquartile range (IQR), horizontal like marks the median, and whiskers extend to farthest data points within 1.5 times the IQR. **(b)** The zero-shot predicted normalized effect on tumor expression using Multicellular Transformer vs the observed effect on tumor across lung cancer cell lines from DepMap data (Tsherniak et al. 2017). The predicted effect from Multicellular Transformer was computed by calculating the change in expression in the sample between perturbed vs control and normalizing it by control expression. **(c)** The zero-shot predicted normalized effect on non-tumor cells vs on tumor cells. The predicted effect was calculated same as in (c). **(d)** The predicted gene knock-out effect on tumor and non-tumor cells on lung cancer (Salcher et al. 2022) and glioblastoma (Ruiz-Moreno et al. 2022) across multiple validation samples on a set of genes specific for lung cancer, glioblastoma and housekeeping genes. The dot size represents the mean expression in the sample, ensuring the gene is expressed in both tumor and non-tumor cells. The dot color signifies the change in relative cell type abundance between the reconstructed expression with vs without the knock-out. The knock-out effect was calculated by zeroing out the gene expression at input and comparing the log2fc in cell type abundance compared to the baseline sample without the gene knock-out. The negative enrichment in reconstructed tumor cell expression compared to non-tumor cells shows how MCT captures tissue-specific tumor-selective dependencies.

#### Predicting the effect of genetic perturbation on cell types

Moving beyond individual gene-gene interactions, we next asked whether MCT could predict the effects of perturbations on different cell types within a population. Because MCT takes multicellular samples as input, it can estimate population-level consequences that cell-line-based approaches do not capture.

To quantify this effect, we compared reconstructed normalized expression aggregated at the cell-type level with and without the knockout (Methods). We benchmarked the predictions against *DepMap* (Tsherniak et al. 2017), which provides CRISPR knockout viability readouts across cell lines.

Using the lung cancer fine-tuned model, we performed *in silico* knockouts across lung cancer samples (Methods) and compared the strongest predicted tumor effects with *DepMap*. MCT showed a 30% overlap with top *DepMap* hits (Supp. Fig. S15b). For the *SW 1573* cell line, the Pearson correlation between MCT and *DepMap* was 0.33 ( *p*-value < 1 × 10^−10^; Supp. Fig. S7b). Many shared high-effect genes were ribosomal proteins, consistent with their general essentiality. To prioritize potentially safer targets, we compared predicted effects on tumor and non-tumor cells (Methods), identifying genes with stronger tumor-specific effects and substantially less enrichment for ribosomal proteins (Fig. 4c).

Because *DepMap* includes hundreds of cell lines, these correlations can also help identify experimental models whose dependencies most closely match the multicellular context captured by MCT (Supp. Fig. S15d); this could potentially help inform disease model design – a critical step in any pre-clinical validation.

We then perturbed disease-specific genes in lung cancer (Salcher et al. 2022) and glioblastoma (Ruiz-Moreno et al. 2022) patient samples and compared the resulting changes in tumor and non-tumor cell populations. Tumor-specific genes consistently reduced tumor populations more than other cell types, whereas house-keeping genes had little selective effect despite high expression (Fig. 4d). Some dependencies were shared across brain and lung tumors, but the effects were strongest for tissue-matched genes, showing that MCT can distinguish context-specific vulnerabilities from broadly essential genes.

### 2.5. Multicellular Transformer recovers spatial co-localization patterns from scRNA-seq data

To determine if MCT captures spatial patterns from scRNA-seq data alone, we tested whether genes proximal in embedding space co-localize in spatial transcriptomics data. We extracted high-similarity gene pairs from MCT embeddings trained on lung cancer (Salcher et al. 2022) and glioblastoma (Ruiz-Moreno et al. 2022) atlases, then validated them using matched Xenium and Visium datasets excluded from training. Gene pairs with high cosine similarity exhibited significant spatial co-localization, as measured by bivariate Moran’s I (Fig. 5a-b; Supp. Fig. S16a-b). Furthermore, the correlation between embedding similarity and spatial co-localization strengthened in higher co-localization percentiles (Fig. 5c), whereas the weaker correlation at lower percentiles reflects the near-zero Moran’s I values for most pairs (Supp. Fig. S16e). These findings show that MCT captures gene interaction patterns consistent with physical co-localization, enabling the recovery of spatially-informed signals from standard scRNA-seq data.

**FIGURE 5.**
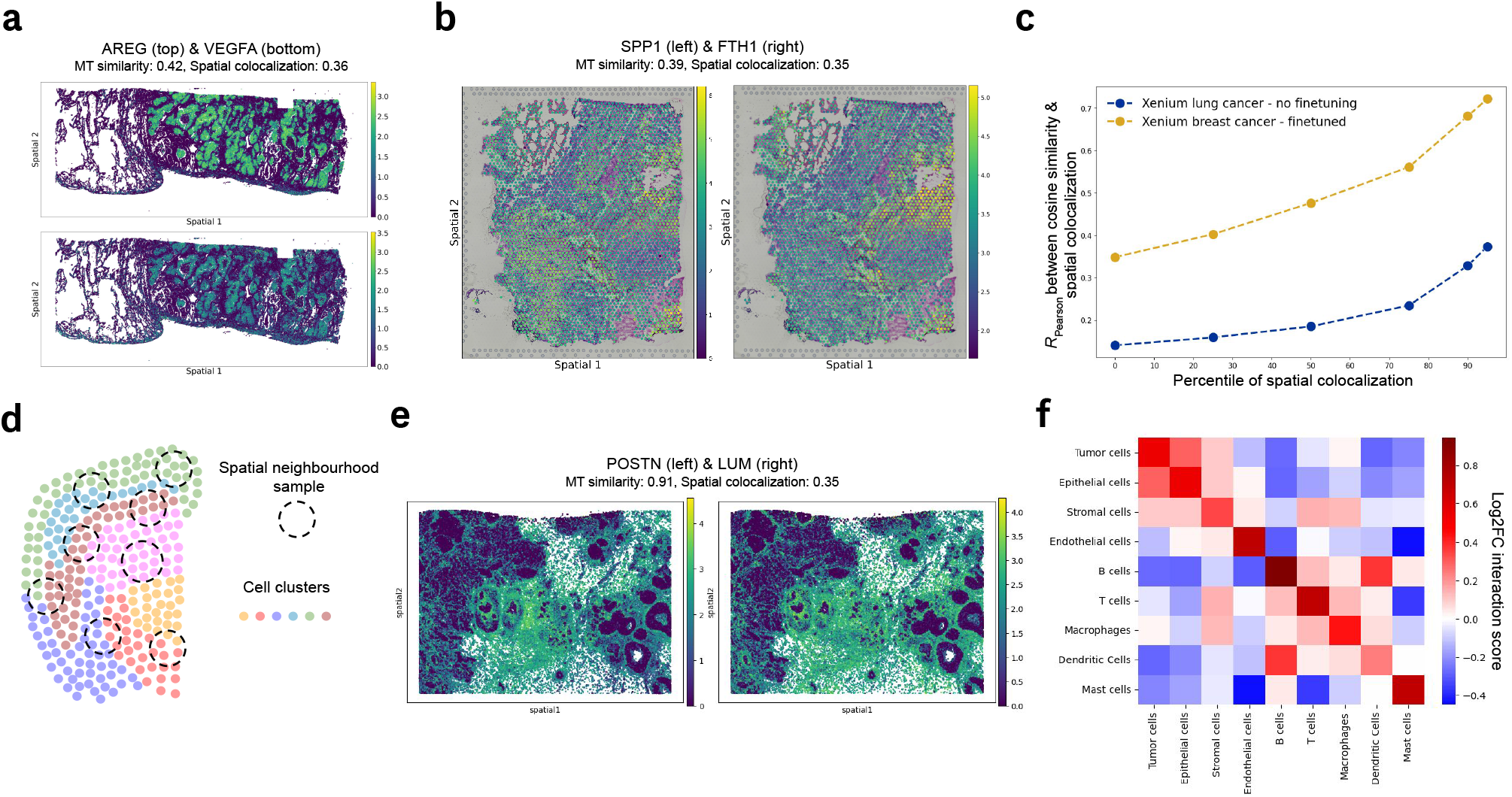
Spatial patterns across datasets recovered by the Multicellular Transformer. **(a)** Spatial co-localization of *AREG* & *VEGFA* genes in Xenium human lung cancer dataset, recovered by the Multicellular Transformer in a zero-shot fashion by assessing genes with highest cosine similarity in gene embeddings for the *AREG* gene. The spatial co-localization was measured by computing bivariate Moran’s I between two genes. **(b)** Spatial co-localization of *SPP1* & *FTH1* genes in Visium glioblastoma dataset, recovered by the Multicellular Transformer in a zero-shot fashion by assessing genes with highest cosine similarity in gene embeddings for the *SPP1* gene. The spatial co-localization was measured by computing bivariate Moran’s I between two genes. **(c)** The correlation between spatial co-localization and the cosine similarity computed using Multicellular’s Transformer gene embeddings across different percentiles of absolute spatial co-localization (the higher the percentile, the higher the absolute spatial co-localization). The correlation was computed by computing spatial co-localization using bivariate Moran’s I, and cosine similarity using gene embeddings from Multicellular Transformer. **(d)** A visualization showing spatial samples used for fine-tuning the Multicellular Transformer model on spatial transcriptomics data. A spatial sample is defined by taking the cells within a predefined radius of a cell of interest. **(e)** Spatial co-localization of *POSTN* & *LUM* genes in Xenium breast cancer dataset, recovered by the Multicellular Transformer fine-tuned on the dataset by assessing genes with highest cosine similarity in the gene embedding space. **(f)** Interaction matrix between cell types computed using the cosine similarity between Multicellular Transformer’s gene embeddings of marker genes. The Multicellular Transformer model used here was fine-tuned on the Xenium human breast cancer dataset and the marker genes were annotated using the same dataset. The interaction score was computed by calculating the *log*2 fold change between the gene pair’s cosine similarity divided by its average cosine similarity across all genes and taking the average across gene pairs for each cell type-cell type pair.

#### Fine-tuning on spatial transcriptomics data

Cell-cell interactions are most prominent within local spatial niches (Morrison and Spradling 2008). While MCT recovers spatial patterns from scRNA-seq alone, we reasoned that performance could be further enhanced by explicitly encoding spatial constraints. We therefore fine-tuned MCT on Xenium breast cancer data, defining samples as cells within a fixed radius of a central cell (Fig. 5d, Methods).

Fine-tuning increased the correlation between embedding similarity and spatial co-localization, recovering more highly co-localized gene pairs than the base model (Fig. 5c,e; Supp. Fig. S16c). This improved resolution is biologically relevant as it enables the identification of interactions between distinct cell types, such as tumor and T cells (Supp. Fig. S16d). Indeed, a cell-type interaction matrix derived from these embeddings revealed distinct immune and stromal compartments that were otherwise obscured by noise in standard co-expression analyses (Fig. 5f; Supp. Fig. S16f). Together, these results demonstrate that while MCT can capture spatial patterns, spatial fine-tuning further refines its ability to recover these.

## 3. Discussion

In this study, we introduced the Multicellular Transformer (MCT), a foundation model that shifts the analytical unit from isolated cells to coordinated multicellular samples. By modeling genes within their joint expression context, MCT aims to link gene-level regulation to tissue-scale behavior, reflecting the hierarchical tissue organization where cellular states are influenced by their surrounding signaling environments.

Our findings suggest that contextual embeddings may serve as a useful tool for biomarker discovery, as evidenced by the disease-specific gene rewiring observed in SLE and COVID-19. Furthermore, the model’s ability to prioritize cell-population-level dependencies offers a potential framework for clinically-relevant tasks, such as *in silico* target prioritization and informing the selection of pre-clinical models. These capabilities are supported by the model’s recovery of ligand-receptor interactions and spatially-informed signals, suggesting that MCT captures meaningful interaction patterns even in dissociated scRNA-seq data, although biases in public dataset coverage, protocols, and annotations may affect generalization to underrepresented contexts.

MCT provides a scalable framework for exploring multicellular biology at a resolution that captures tissue-scale complexity. By moving beyond isolated cells, this approach may facilitate the discovery of coordinated gene programs and therapeutic vulnerabilities. While MCT represents a valuable contribution, we acknowledge certain limitations that offer promising avenues for future research. For example, further work is needed to derive mechanistic insights into *how* gene entities interact rather than simply determining *whether* they interact. Another challenge lies in designing more direct methods to pinpoint the specific cell subsets between which interactions occur, as our current framework relies on post-hoc explainability techniques (e.g., Integrated Gradients) to obtain these insights. Finally, although our fixed input allocation for predefined cell types ensures structured cross-cell learning, it currently constrains zero-shot generalization to entirely novel, unannotated cell types and requires handling variations in cell type proportions through sampling strategies. Nevertheless, because MCT was pretrained across more than 300 harmonized cell types, it natively covers the primary cell populations of most standard biological samples. Future work will focus on addressing these challenges, expanding the pretraining corpus, and integrating protein-level or multimodal data. Despite there being areas of improvement, we believe that MCT offers a new lens for modeling collective cellular behavior and the drivers of human disease.

## 4. Data Availability

All datasets used in this study are publicly available. Details and accession information for the datasets used for pretraining are provided in Supplementary Table S1. Details of all datasets used in downstream analyses, together with the corresponding preprocessing procedures, are provided in the Methods. The pretraining corpus was stored, accessed, and sampled using the CellArr package (https://github.com/BiocPy/cellarr).

## 5. Code Availability

The code used to pretrain Multicellular Transformer and reproduce the downstream experiments is publicly available at https://github.com/almaan/mct. The repository also includes tutorials and documentation for using Multicellular Transformer.

## 6. Author Contribution

M.W. led the project, developed the methodology, and performed the majority of the analyses and experiments. A.A. co-supervised the project, provided scientific feedback on the methodology, contributed to the analyses, and contributed substantially to writing and editing the manuscript. K.H.H. contributed to data analysis and scientific discussions. T.B. provided scientific feedback and guidance. A.B. conceived and supervised the project and provided scientific direction and guidance. All authors reviewed and approved the final manuscript.

