## Supplementary Material for "Learning gene co-regulation across cells with Multicellular Transformer"

**3 Discussion** **13**

**4 Data Availability** **13**

**5 Code Availability** **14**

**6 Author Contribution** **14**

**7 Competing Interests** **14**

**A Methods** **1**

**B Supplementary Figures & Tables** **14**

**C Sensitivity of contextual embeddings to cell ordering and resampling** **31**

**D Scaling Laws** **33**

### Appendix A. Methods

#### A.1. Model architecture & training

Multicellular Transformer takes as input a set of of cells originating from the same sample, allowing it to model the interactions between genes across cells. Therefore, in contrary to a number of existing single-cell foundational models (Cui et al. 2024; Theodoris et al. 2023; Hao et al. 2024; Heimberg et al. 2023), a single input example is not a single cell, but a cell-by-gene matrix $X \in \mathbb{R}^{M \times N}$ originating from the same sample, where $M$ is the number of cells and $N$ is the number of genes. Here, the sample means cells coming from the same donor, tissue and sequencing batch. Therefore, the cells which may interact with each other. This matrix is then used for modeling the contextual interactions between genes in a multicellular environment.

##### A.1.1. Multicellular gene embeddings

The multicellular gene embeddings, used as input to the transformer, consist of two parts: the gene expression latent vector and the gene identity embeddings. The first represents the context-dependent activity of the gene across cells in the sample, while the second represents the context-independent function of the gene, which is the same across all samples. The two are computed separately and then combined by summation.

**Gene expression embeddings**. Given an expression matrix $X \in \mathbb{R}^{N \times M}$, where $N$ is the number of genes in a sample and $M$ is the number of cells, each element $X_{i,j} \in \mathbb{R}^{+}$ represents the normalized expression value (after $log1p$ transformation) of gene $g_i$ in cell $c_j$.

For each gene $g_i$, we define its expression vector $\mathbf{x}_{g_i} \in \mathbb{R}^{M}$ as the $i$-th row of $X$:

$$\mathbf{x}_{g_i} = [\, X_{i,1},\ X_{i,2},\ \ldots,\ X_{i,M} \,]$$

This vector represents the expression of gene $g_i$ across all $M$ cells in the sample.

The feed-forward neural network (FFNN) $MLP_e$ is a 2-layer multilayer perceptron (MLP) which transforms the gene expression vector across cells into a latent vector of dimension $D$:

$$\text{GEXEmb}_{g_i} = MLP_e(\mathbf{x}_{g_i})$$

The $\text{MLP}_e$ is shared across all genes to prevent overfitting, therefore, it is not gene-specific.

Importantly, the input units of $MLP_e$ of dimension $M$ are associated with certain cell types during training. Specifically, given a set of cell types $\{t_1, t_2, \ldots, t_T\}$, where $T \leq M$, the input vector $\mathbf{x}_{g_i}$ can be partitioned according to these cell types:

$$\mathbf{x}_{g_i} = [\, \mathbf{x}_{g_i}^{t_1},\ \mathbf{x}_{g_i}^{t_2},\ \ldots,\ \mathbf{x}_{g_i}^{t_T} \,]$$

where $\mathbf{x}_{g_i}^{t_k}$ contains the expression values of gene $g_i$ in cells of type $t_k$, such as *CD4 T cell*. Fixing input units to certain cell types allows the model to learn cell type-specific patterns, instead of just learning average gene loadings, and is crucial for the model to learn meaningful interactions.

It is also possible to use a pseudobulked input vector as input to the model. In this case, we aggregate the expression values of each gene across all cells of the same cell type, resulting in a reduced expression vector that captures the total expression per cell type. Formally, for each gene $g_i$, we compute a pseudobulked expression vector $\mathbf{p}_{g_i} \in \mathbb{R}^T$, where $T$ is the number of cell types ($T \leq M$). Each element $p_{g_i}^{t_k}$ in $\mathbf{p}_{g_i}$ represents the sum of normalized expression values of gene $g_i$ across all cells belonging to the cell type $t_k$:

$$p_{g_i}^{t_k} = \sum_{j \in C_{t_k}} X_{i,j}$$

where $C_{t_k}$ is the set of cell indices corresponding to cell type $t_k$. In this pseudobulked approach, the input dimension to the gene expression network $\text{MLP}_e$ becomes $T$ instead of $M$, reducing the input size and focusing on aggregated expression patterns per cell type. We have trained models using both the original per-cell expression vectors $\mathbf{x}_{g_i} \in \mathbb{R}^M$ and the pseudobulked expression vectors $\mathbf{p}_{g_i} \in \mathbb{R}^T$, with both approaches performing competitively. Model weights for both versions are made available.

We have also experimented with training the model without utilizing cell type labels, effectively treating all cells as a single group. The resulting model significantly underperformed compared to the one using cell type labels, highlighting the importance of incorporating cell type information.

Finally, during pretraining, we used cell type labels to guide the partitioning and aggregation of expression vectors. However, it is also possible to use any mutually exclusive cell labels, such as clusters derived from unsupervised methods like Leiden clustering. We demonstrated this when fine-tuning the Multicellular Transformer on spatial transcriptomics data, where we utilized labels from Leiden clustering performed on the dataset.

**Gene identity embeddings**. Given a predefined set of genes $G$, we assign each gene $g_i$ a unique identifier, which forms a vocabulary of tokens used in the Multicellular Transformer. This allows us to harmonize different sets of genes across different studies, rather than taking the intersection of genes across them. We use these identifiers to transform a gene into a multidimensional vector of dimension $D$, using a gene embedding layer function $\text{emb}_g$, which is learned during pretraining. The output of the gene embedding layer $\text{emb}_g$ is the same size as the output of the gene expression $\text{MLP}_e$.

Combining all of the above, for a gene $g_i$ with expression vector $\mathbf{x}_{g_i} \in \mathbb{R}^M$, the multicellular gene embedding is defined as:

$$\text{Embedding}_{g_i} = \text{emb}_g(g_i) + \text{MLP}_e(\mathbf{x}_{g_i})$$

Across a set of genes present in the sample, the input to the transformer is:

$$\text{Input} = [\, \text{Embedding}_{g_1},\ \text{Embedding}_{g_2},\ \ldots,\ \text{Embedding}_{g_L} \,],$$

where $L$ denotes a predefined maximum input length. This setup allows us to: (a) account for multiple cells in the environment and the interactions between them, (b) consider the context-dependent gene expression of the genes present in the sample, (c) represent gene function independent of expression across samples.

##### A.1.2. Transformer block

We leverage the transformer (Vaswani et al. 2017) architecture to compute contextual representations of multicellular gene embeddings. Given the expression matrix $X \in \mathbb{R}^{M \times N}$, we first compute the multicellular gene embeddings for each gene $g_i$ as previously defined:

$$\text{Embedding}_{g_i} = \text{emb}_g(g_i) + \text{MLP}_e(\mathbf{x}_{g_i})$$

where $\mathbf{x}_{g_i} \in \mathbb{R}^M$ is the expression vector of gene $g_i$ across all $M$ cells in the sample.

We collect these embeddings into a sequence:

$$\mathbf{E} = [\,\text{Embedding}_{g_1},\ \text{Embedding}_{g_2},\ \ldots,\ \text{Embedding}_{g_L}\,]$$

where $L$ is the number of genes considered (up to a predefined maximum input length).

We then compute their contextual representations $\mathbf{H} = [\,\mathbf{h}_1,\ \mathbf{h}_2,\ \ldots,\ \mathbf{h}_L\,]$ by passing them through multiple transformer layers:

$$\mathbf{H} = \text{Transformer}(\mathbf{E})$$

To improve the efficiency of the self-attention mechanism, we use Flash Attention (Dao et al. 2022). Additionally, we include special tokens [CLS] and [SEP] to mark the beginning and end of the sequence.

##### A.1.3. Training objective

The model is optimized with two objectives: (1) a masked modeling objective (Devlin 2018), and (2) a gene expression reconstruction objective.

In the first objective, we randomly mask 15% of the genes in the input sequence. Specifically, we design our own masking strategy, where out of the 15% of masked genes:

- 70% are masked gene identity embeddings: for gene $g_i$, we replace its gene identity embedding $\text{emb}_g(g_i)$ with a special token embedding $[MASK]$.
- 10% are masked gene expression embeddings, where we replace the gene expression embedding with $[MASK]$ vector,
- 10% where random gene identity embedding is swapped with the true gene identity embedding,
- 10% where neither gene identity nor expression is masked, leaving the embeddings unchanged.

This strategy is inspired by the masking strategy in BERT (Devlin 2018) and allows the model to learn robust and context-aware representations by predicting masked gene identities and expressions. By varying the masking types—including masking gene identities, gene expressions, swapping gene identities, and leaving some unchanged—the model is encouraged to leverage both gene identity and expression information. This helps the model capture complex dependencies among genes and understand the impact of gene expression patterns across different cells, thereby enhancing its ability to model gene interactions in a multicellular environment. The gene probabilities for each masked gene are computed by passing the output of a transformer $\mathbf{h}_i \in \mathbb{R}^D$ of a masked gene $g_i$ through a classifier head which is weight matrix $W \in \mathbb{R}^{D \times G}$, where $D$ is the embedding dimension and $G$ is the total number of genes in the pretraining corpus. $W$ is learned over the course of pretraining and used for analysis on downstream tasks. The masked modeling loss $\mathcal{L}_{\text{mask}}$ is defined as:

$$\mathcal{L}_{\text{mask}} = -\frac{1}{|\mathcal{M}|} \sum_{i \in \mathcal{M}} \log P(g_i \mid \mathbf{E}_{\text{masked}})$$

where $\mathcal{M}$ is the set of positions of masked gene identities, $g_i$ is the true gene identity at position $i$, and $\mathbf{E}_{\text{masked}}$ represents the input embeddings with applied masking.

For the gene expression reconstruction objective, we use the last hidden state from the transformer block to reconstruct the input normalized gene expression for each gene. This is performed for all genes in a sample, including genes with masked gene identity as well as masked gene expression. To reconstruct the gene expression, we learn a 2-layer MLP, $MLP_r$, which, given a $D$-dimensional vector, projects it into an $M$-dimensional vector equal to the input gene expression vector:

$$\hat{\mathbf{x}}_{g_i} = \text{MLP}_r(\mathbf{h}_i)$$

where $\mathbf{h}_i \in \mathbb{R}^D$ is the hidden representation of gene $g_i$ from the transformer, and $\hat{\mathbf{x}}_{g_i} \in \mathbb{R}^M$ is the reconstructed gene expression vector. The reconstruction loss $\mathcal{L}_{\text{recon}}$ is defined as the mean squared error between the reconstructed and true gene expression vectors:

$$\mathcal{L}_{\text{recon}} = \frac{1}{L} \sum_{i=1}^{L} \left\| \hat{\mathbf{x}}_{g_i} - \mathbf{x}_{g_i} \right\|^2$$

where $L$ is the total number of genes in the input sequence.

The final loss function consists of the two losses described above, weighted by $\alpha$ (masked cross-entropy loss) and $1 - \alpha$ (reconstruction loss):

$$\mathcal{L} = \alpha \mathcal{L}_{\text{mask}} + (1 - \alpha) \mathcal{L}_{\text{recon}}$$

we experimented with different values of $\alpha$ and used $\alpha = 0.99$ in the best performing pretraining run. Large value of $\alpha$ is due to the reconstruction loss having significantly larger absolute values. By optimizing this combined loss function, the model learns to predict masked gene identities and accurately reconstruct gene expression profiles, capturing intricate patterns and dependencies within the multicellular context. We have found that the masked learning objective is the most important to learning gene-gene interactions which we are interested in. However, the gene expression reconstruction improves the stability of training and allows us to use it for downstream tasks, such as *in silico* perturbation effect prediction at cell population level.

##### A.1.4. Training details

The transformer (Vaswani et al. 2017) block consists of 12 transformer layers with self-attention. The gene expression embedding $MLP_e$ and gene expression reconstruction $MLP_r$ are both 2-layer MLPs, interleaved with *ReLU* activation function (Agarap 2018), layer normalization (Ba 2016) and dropout (Srivastava et al. 2014) equal to 0.2. The gene identity embedding is an embedding layer $emb_g \in \mathbb{R}^{G \times M}$. All parameters are learned from scratch. Total nr of parameters equals ≈ 100M. The model in implemented in PyTorch (Paszke et al. 2019).

We have performed random split on sample-level, keeping 0.95 for training and 0.05 for validation. We did not include a test set as we do not have a downstream task which directly matches our pretraining scenario. During pretraining, we set the number of cells in an input example of the model to $1,024$ and filter genes which are expressed (normalized expression > 0) in less than 1% of the sampled cells. The maximum context size, which here is the maximum nr of genes in a sample, was set to $12,000$.

We have trained the model with the learning rate 0.00015, *Adam* optimizer (Kingma 2014) with weight decay and a linear warmup of $10,000$ steps. We have performed validation every $10,000$ steps, monitoring the validation loss. We used the batch size of 4 while accumulating gradients for 4 batches before performing a backward pass. The maximum nr of training steps has been set to 500,000. The training has been performed on a single NVIDIA A100 80GB GPU for approximately 14 days.

#### A.2. Data preprocessing & training data augmentation

**scRNA-seq preprocessing**. Multicellular Transformer takes as input a gene-by-cell expression matrix $X \in \mathbb{R}^{N \times M}$, where $N$ is the number of genes and $M$ is the number of cells, each element $X_{i,j} \in \mathbb{R}^+$ is the normalized expression value. The normalization is performed by normalizing each cell by total counts over all genes, so that every cell has the same total count after normalization. We set the target counts sum to $10,000$. Then, $log1p$ normalization is applied. The preprocessing was performed using `scanpy` package (Wolf, Angerer, and Theis 2018).

**Cell type harmonization**. Multicellular Transformer learns interactions between genes across cells, therefore, in contrary to existing single-cell foundational models, the input is not the expression of a gene in a single-cell, but the expression of a gene across distinct cells. The cells within and across samples differ, and we cannot rely on the same genes as features for every sample, therefore, to ensure consistency, the input units of the gene expression embedding, $MLP_e$ are reserved for certain cell types during pretraining (see *Gene expression embeddings* section above). However, cell types across datasets often differ, therefore, we harmonize more than 300 cell types present in the datasets used for pretraining to $X$ cell types. We recalculate the relative number of cells for each cell type, and reserve a number of input units in the $MLP_e$ proportional to the relative count of the cell type. Finally, we remove the cells belong to cell types with less than 1% of relative counts.

**Training data augmentation**. An input example to the Multicellular Transformer is a sample, and our pretraining corpus has around 4,200 distinct samples, which is a low number for training a model of this size, even considering the large diversity of the samples. Moreover, Multicellular Transformers requires $M$ cells as input in every sample, which is used as input to gene expression embedding MLP, $MLP_e$. Therefore, we employ a training data augmentation technique by iteratively sampling $M$ cells from each sample. This prevents the model from simply remembering, thus overfitting to each sample, as every time the set of sampled cells is different, and many samples have more than $10,000$ cells. We sample the cells from each sample using stratified sampling, where the class is a cell type label. This ensures consistency over various samples. The cell type counts used for stratified sampling are precomputed prior to training. In some samples, there may not be enough of cell types of a particular class, for example, sample $A$ has only 70 *Macrophages*, while the $MLP_e$ required 80. In this case, we pad the input with zeros. The training data augmentation played a crucial role in our pretraining, with the model without it performing significantly worse.

**Gene filtering**. Across cells, the majority of the genes will have scRNA-seq counts equal to 0. This is due to the presence of dropout in the transcriptomic data. Existing single-cell foundational models, such as scGPT drop the genes with 0 counts, as this would make predicting gene identity challenging, causing the model to fail to converge, as the huge majority of genes will have the same expression level, equal to 0. This is a severe limitation as it makes the model only learn the interactions between the genes with expression higher than 0, which is usually around 1,000 genes.

Multicellular Transformer takes as input a set of cells and their corresponding scRNA-seq counts across genes. Therefore, instead of filtering genes at a single-cell level, we filter genes at the sample-level. Specifically, during pretraining we filter out genes which are expressed (normalized expression > 0) in less than 1% of cells. This allows us to significantly increase our gene context size and take as input $\approx 12,000$ genes, rather than $\approx 1,000$ genes in a single-cell scenario. Our results suggest, that the context size plays a crucial role on downstream tasks, such as gene set extraction and receptor-ligand identification. We have also experimented with more or less stringent filtering and found that more stringent filtering leads to a stable training, however, worse results on downstream tasks, while the less stringent filtering leads to less stable training and often similar or worse performance on downstream tasks. The context size used during our pretraining was set to $12,000$ genes.

**Sampling robustness analysis**. During training and validation, Multicellular Transformer samples $M$ cells from every sample, where $M$ is a parameter set prior to training ($M = 1,024$ during pretraining). This serves as a training data augmentation technique, as the number of samples is relatively low, but each sample contains a different number of cells. Therefore, by sampling from each sample, we increase the effective training data size and introduce variability, helping the model generalize better across distinct cell populations. However, by sampling from each sample, we may also introduce noise if the result of sampling from the same sample varies too much. To assess it, we have run sampling on 100 randomly selected samples from the validation set of the pretraining corpus, sampling each sample 5 times. We have then extracted 50 nearest neighbors for each gene across samples, and evaluated the overlap between the gene neighborhoods within the same sample as well as across samples. The overlap within a sample across genes is signicantly higher than the one across samples, achieving the median of 0.2 Jaccard similarity, compared to 0.08 between distinct samples. Therefore, showing that MT can robustly account for the variability within the same sample, maintaining relatively stable gene neighborhoods compared to the gene neighborhoods across distinct samples.

#### A.3. Downstream tasks & analysis

##### A.3.1. Receptor-Ligand identification

To analyse the interactions between receptor and ligand genes learned over the course of pretraining, we extracted the pretrained gene embedding matrix from the masked learning classifier, resulting in a $X \in \mathbb{R}^{\mathbb{G} \times \mathbb{D}}$ matrix, where $G$ is the total number of genes present in the pretraining corpus and $D$ is the dimensionality. Using a predefined set of true and false receptor-ligand pairs, we have computed the cosine similarity value for each pair and employed it to calculate the area under the ROC curve (AUC).

**Dataset**. To evaluate the performance on the receptor-ligand identification task, we have used a set of known consensus receptor-ligand pairs from *LIANA* (Dimitrov et al. 2024). Importantly, not all of these pairs have been experimentally validated and may not be physically present on the cell surface for the receptors and secreted for the ligands. However, they represent a set of consensus pairs across multiple receptor-ligand sources and have been shown to play a crucial role in cell-cell communication. Therefore, we use it for evaluation.

To establish negative pairs, which would complement the set of positive receptor-ligand pairs described above, we randomly created negatives using genes present in all the benchmarks in a ratio 1:10 for positives to negatives. Importantly, to avoid only including easy negatives with no overlap with the positive pairs, half of the negatives were created by using a receptor present in the true positives. This is to avoid a situation, where the positives and negatives have no overlap in the genes, making it potentially significantly easier for the model and inflating the results. In our case, we wanted to evaluate whether the model can identify the correct ligands for a receptor from a large pool of potential ligands. Finally, our setup does not account for false negatives, which may arise as the space of receptor-ligand remains unexplored. However, the statistical chances of it are small and the evaluation is the same for all models, therefore making it comparable.

For fair comparison, we only considered the receptor-ligand pair which were present in all benchmarks used for the analysis. Additionally, as our analysis was transcriptomics-based, we excluded pairs where either a receptor or ligand are a protein complex created from more than one gene product.

**Benchmarks**. We followed the same approach for extracting gene embeddings for scGPT and CellPLM as in the gene set extraction analysis. We then computed the cosine similarity for all pairs using both benchmarks and used this score for computing metrics. For co-expression, we followed the same steps as in the section on gene set extraction analysis to get a matrix $E \in \mathbb{R}^{GxS}$, where $S$ represents the number of samples in the pretraining corpus, and $G$ total number of genes. We then computed the *Pearson correlation* coefficient for all pairs of genes in the dataset and used the resulting coefficient for computing metrics. Finally, for context size impact analysis. We have used Multicellular Transformer models pretrained with different context size.

**Integrating multicellular and protein-level information**. To complement MCT with protein-level context, we employed the ESMC protein language model (ESM Team 2024), a state-of-the-art approach for learning protein representations. Using UniProt, we mapped the *Ensembl* gene IDs in the LIANA consensus receptor–ligand set (Dimitrov et al. 2024) to their corresponding UniProt protein IDs and retrieved the associated amino-acid sequences. Each sequence was embedded with the open-source 300 M-parameter ESMC model, averaging the residue embeddings to obtain a single vector for every protein. We then computed the cosine similarity between receptor and ligand vectors to produce a scalar interaction score. This ESMC score was concatenated with the MCT score to form a two-dimensional feature vector, which served as input to an XGBoost classifier. We performed five-fold cross-validation and report the mean performance across folds.

**Identifying protein complexes using contextual gene embeddings.** We followed the same approach to compute cosine similarities between contextual gene embeddings as in the paragraph above. However, we did not limit ourselves to receptor-ligand pairs and compute cosine similarity across both disease and healthy together. We focused on cytokines, as they play key part in immune system signaling. We then computed cosine similarities across all cytokines and filtered pairs which appeared in less than 3 samples. Finally, we only considered pairs where one of the pairs was secreted from the cell and the other was a transmembrane protein. We have retrieved this information from UniProt (UniProtConsortium 2019). The top 70 pairs were screened using AlphaFold3 (Abramson et al. 2024) server.

**Analysing receptor-ligand interactions in healthy vs disease.** We used the systemic lupus erythematosus (SLE) study (Perez et al. 2022), and ran Multicellular Transformer across all samples. For each sample, we have computed the cosine similarity using contextual embedding of receptor-ligand pairs from LIANA consensus set (Dimitrov et al. 2024) present in the sample, keeping track of whether the sample has been from disease or healthy patient. After collecting this data, we have excluded the receptor-ligand pairs which appeared less than 3 times in either healthy or disease to remove pairs for which the resulting analysis would not be interpretable due to low sample size. Then, we calculated the relative change in the cosine similarity value in disease vs healthy. Let $X$ represent the cosine similarity of the receptor-ligand pair in disease samples, and $Y$ represent the cosine similarity in healthy samples. Then, the relative fold change in cosine similarity $\Delta_{\mathrm{rel}}$ is given by $\Delta_{\mathrm{rel}} = \frac{X-Y}{Y}$.

##### A.3.2. Gene set extraction & analysis

Similarly as with receptor-ligand analysis, to analyse the interactions between genes learned by the model, we have extracted the pretrained gene embedding matrix from the masked learning classifier, resulting in a $X \in \mathbb{R}^{\mathbb{G}\times\mathbb{D}}$ matrix, where $G$ is the total number of genes present in the pretraining corpus and $D$ is the dimensionality. We have also experimented with using the gene identity embeddings, but found the former to perform better across different tasks. We have followed the setup for gene set extraction used in scGPT (Cui et al. 2024) for comparability. The gene embeddings were used to compute the gene-gene similarity matrix. We have then used the gene-gene similarity matrix to cluster genes into gene sets by applying Leiden clustering (Traag, Waltman, and Van Eck 2019). The gene sets are mutually exclusive, meaning if a gene was present in gene set A, it could not be present in any other gene set. We have run the Leiden clustering across different resolutions, setting minimum resolution at 10, as lower resolutions led to very large gene sets, which were not interpretable (Supp. Fig. S6b). Following clustering, we have excluded the gene sets with fewer then 4 genes.

**Cell-cell communication pathways enrichment**. To assess whether the extracted gene sets were enriched in cell-cell communication related pathways, we selected the pathways from `GO` (Ashburner et al. 2000) and `Reactome` (Fabregat et al. 2018) and filtered them for gene sets which included either of predefined keywords: `[receptor, ligand, signal transduction, cell signaling, signaling pathway, signaling pathway, communication]`. The extracted pathways were then manually verified. This left us with 451 CCC related GO pathways and 137 CCC related Reactome pathways. We used `GSEApy` (Fang, Liu, and Peltz 2023) to identify pathways enriched in gene sets, and kept only the significantly enriched pathways. We performed enrichment using both `GO` and `Reactome` libraries.

We used the highly variable genes from the tumor microenvironment atlas, enriched with receptor and ligand genes, setting the total number of genes to 8,000. We only considered the genes which were present in all benchmarked methods. The tumor microenvironment atlas (Guimarães et al. 2024) was selected as (1) the multicellular interactions are particularly important in cancer, (2) none of the methods have been trained on the dataset, therefore the setup was zero-shot.

**Benchmarks**. For *scGPT* (Cui et al. 2024) and *CellPLM* (Wen et al. 2023), we used the latest recommended checkpoints to extract gene embeddings (whole human, pretrained on 33M human cells for scGPT and checkpoint `20230926_85M` for CellPLM). We then followed the same framework for gene set extraction and analysis as for the Multicellular Transformer described above. For co-expression, we have aggregated the normalized expression across samples, summing over the expression over cells for each gene in the sample. Therefore, focusing on the intercellular, rather than intracellular expression achieved by calculating the co-expression at cell-level. This results in a matrix $E \in \mathbb{R}^{GxS}$, where $S$ represents the number of samples in the pretraining corpus, and $G$ total number of genes. We then computed the PCA (Abdi and Williams 2010) on matrix $E$, resulting in a matrix $E_{PCA} \in \mathbb{R}^{GxP}$, where $P$ is the number of principal components (here, 50). The $E_{PCA}$ matrix was used to cluster genes with Leiden clustering, same as for the methods above.

We provide a jupyter notebook outlining how to extract gene sets given multicellular gene embeddings.

##### A.3.3. Context-aware gene embeddings analysis

For context-aware gene embeddings analysis, we used the output representations from the final transformer layer. We wanted to identify condition-specific genes, which we defined as genes with consistent neighborhood within a condition, while simultaneously having a different neighborhood across conditions.

To calculate this, we retrieved the 50 nearest neighbors for each gene across all samples from three datasets: 1) immune system across organs (Suo et al. 2022), 2) systemic lupus erythematosus (Perez et al. 2022), and 3) COVID-19 (Stephenson et al. 2021). We then computed the mean overlap across neighborhoods within a condition by calculating the Jaccard similarity between the neighborhoods of each gene across samples with the same condition (e.g., from the same tissue or disease) using the 20 nearest neighbors.

For each gene $g$, let $N_g^{(s,c)}$ denote the set of 20 nearest neighbors in sample $s$ with condition $c$. The mean within-condition Jaccard similarity $J_{\text{within}}^{(c)}(g)$ for gene $g$ in condition $c$ is computed as follows:

$$J_{\text{within}}^{(c)}(g) = \frac{1}{|S_c| \cdot (|S_c| - 1)} \sum_{\substack{s,s' \in S_c \\ s \neq s'}} \frac{|N_g^{(s,c)} \cap N_g^{(s',c)}|}{|N_g^{(s,c)} \cup N_g^{(s',c)}|}$$

where $S_c$ is the set of samples under condition $c$.

Next, we computed the Jaccard similarity $J_{\text{across}}^{(c_1,c_2)}(g)$ between each pair of conditions $c_1$ and $c_2$ for gene $g$:

$$J_{\text{across}}^{(c_1,c_2)}(g) = \frac{|N_g^{(c_1)} \cap N_g^{(c_2)}|}{|N_g^{(c_1)} \cup N_g^{(c_2)}|}$$

where $N_g^{(c)}$ represents the most common 40 neighbors of gene $g$ within condition $c$.

Calculating the overlap between samples for each gene is computationally expensive, requiring $G \times |S| \times |S|$ Jaccard similarity calculations, where $G$ is the number of genes and $S$ is the number of samples per condition. To reduce this, we extracted the most common 40 neighbors for each gene within a condition and used these to calculate the overlap across conditions.

We excluded conditions with too few samples to allow for meaningful analysis. Finally, we calculated the fold change between the mean Jaccard similarity within a condition vs. across conditions:

$$\Delta J(g) = \frac{J_{\text{within}}^{(c)}(g)}{J_{\text{across}}(g)}$$

We used this fold change $\Delta J(g)$ to identify condition-specific genes.

##### A.3.4. Sample-level clustering

To extract sample-level embeddings for clustering. We ran Multicellular Transformer in an inference mode and computed sample-level embeddings by averaging out all representations from the final layer of transformer. This results in a matrix $X \in \mathbb{R}^{S \times D}$, where $S$ is the number of samples and $D$ is the dimensionality of the representations. We then ran UMAP (McInnes, Healy, and Melville 2018) on the relevant samples with the number of neighbors varying from 15 to 30 depending on the dataset the samples originated from.

To calculate clustering metrics, we have performed Leiden clustering (Traag, Waltman, and Van Eck 2019) at the resolution of 1.0 and computed the Adjusted Rand Index (ARI) as well as Normalized Mutual Information (NMI). We have also experimented with different levels of Leiden clustering resolution, with the 1.0 consistently showing the best results across methods and datasets.

##### A.3.5. Recovering spatial patterns from scRNA-seq

**Spatial colocalization analysis**. We computed the spatial colocalization by calculating the bivariate Moran's $I$. The bivariate Moran's $I$ is given as:

$$I_{XY} = \frac{N \sum_i \sum_j w_{ij}(X_i - \bar{X})(Y_j - \bar{Y})}{\sum_i (X_i - \bar{X})^2 \sum_i \sum_j w_{ij}}$$

where $N$ is the total number of observations (cells or spots), $X_i$ and $Y_j$ are the values of the two variables $X$ and $Y$ for cells $i$ and $j$, $\bar{X}$ and $\bar{Y}$ are the means of $X$ and $Y$, $w_{ij}$ is the spatial weight between observations $i$ and $j$, where $w_{ij} = 1$ if $i$ and $j$ are neighbors and $w_{ij} = 0$ otherwise.

For the Xenium data, we defined the neighborhood as cells within the radius of 50 µm of the target cell, while for Visium data, we defined it as 6 neighboring spots in the grid. We used the `Squidpy` package (Palla et al. 2022) to perform the analysis. We calculated the bivariate Moran's I for a random subset of 200 genes, meaning we calculated the pairwise Moran' I index for 200 × 200 pairs. This covers the majority of the genes as the gene panels in the Xenium datasets used do not exceed 400 genes after filtering out lowly expressed genes. The cosine similarity has been calculated by extracting the similarity using the contextual gene embeddings between genes across relevant samples. Therefore, for Xenium lung cancer the cosine similarites were computed by running the Multicellular Transformer model in the inference mode across samples from lung cancer atlas (Salcher et al. 2022), and for Xenium breast cancer it was performed by running the Multicellular Transformer fine-tuned on the same dataset across samples in the inference mode. For calculating the correlation between cosine similarity across percentiles, for each percentile, we limited the pairs of genes to values above the percentile and then recalculated the correlation coefficient only for these pairs.

**Fine-tuning on spatial transcriptomics data**. We used the Xenium breast cancer dataset to perform fine-tuning on the spatial transcriptomics data. Due to the lack of annotated cell types, we performed Leiden clustering using default parameters in the `scanpy` package. We then used the resulting annotations and used them as input to the model treating them as cell type labels. Therefore, showing how one can leverage the model without available cell type labels. As a sample, we used the neighboring cells of within a radius of 350 µm of the cell of interest. We then fine-tuned the model using the pretrained objective, leveraging the pretrained gene identity embeddings and transformer block, while learning new $MLP_e$ and $MLP_r$, allowing the model to adapt to the new dataset. We have run the fine-tuning with the same hyperparameters, except the number of cells per sample $M$ being set to 512.

**Cell type-Cell type interactions**. We leveraged the gene annotations from the `10xGenomics` dataset explorer[1] to associate genes with particular cell states. A table with a set of genes per cell type can be found in Supp. Table S2. To calculate the interactions between cell types, we calculated the pairwise similarity between all genes associated with a cell state. Then, to account for the mean cosine similarity of each gene, we divided the resulting cosine similarities by the mean cosine similarities of each gene. Therefore, for genes $g_x$ and $g_y$, the normalized cosine similarity is given as

$$2\frac{g_{xy}}{\bar{g_x} + \bar{g_y}}$$

where $g_{xy}$ represents the cosine similarity between $g_x$ and $g_y$, and $\bar{g_x}$ and $\bar{g_y}$ are the mean cosine similarities

[1] https://www.10xgenomics.com/products/xenium-in-situ/human-breast-dataset-explorer

of $g_x$ and $g_y$ across all genes, respectively. We have then aggregated the results for each cell type-cell type pair, taken a mean of it and applied $log2$ transformation.

We have followed the same approach for calculating cell type-cell type interaction matrix using the spatial colocalization, using spatial colocalization instead of cosine similarity as the measure of similarity.

###### A.3.6. *In silico* perturbation prediction at cell population level

**Calculating gene-gene relationships upon *in silico* genetic perturbations.** We calculated the perturbation effect on other genes by removing the gene from the input genes and evaluating the impact of the gene removal on the predicted gene logits. Specifically, we followed a similar approach to the one proposed in Silva et al. (2024). For a vector of predicted gene logits $b_i$ for a gene $i$ without a perturbed gene and a vector of predicted gene logits $p_i$ with a perturbation, where both logits vectors are of dimension $G$, equal to the total nr of genes, the predicted perturbation effect $pef_i$ is given as:

$$\mathrm{pef}_i = \max\left(\log_2\left(\frac{\mathrm{b}_i}{\mathrm{p}_i}\right)\right).$$

Note that removing the gene from the input is equivalent to zeroing out the gene expression in the Multicellular Transformer setup, as not expressed genes are removed from the input. We ran the *in silico* perturbation across validation samples from the lung cancer atlas, performing the knock-out on each sample where the gene of interest was present, using the Multicellular Transformer model fine-tuned on lung cancer atlas. We then took the mean perturbation effect for true interacting genes and control for each perturbed gene and normalized the results for each knocked-out gene with min max normalization for consistency.

We have used the same dataset as for receptor-ligand task, with the difference that we added negatives following the same approach for the ligands as well. Here the negative pairs served as control. We then selected a subset of the genes to perform the *in silico* gene perturbation.

**Predicting the effect of genetic perturbation across cell types.** Using trained Multicellular Transformer model, we measured the effect of a gene knock-out on a cell type by removing the gene from the input sample and measuring the change in reconstructed gene expression across cell types vs the baseline reconstructed gene expression. Specifically, for a baseline reconstructed normalized expression in cell type $t$, $b_t$, and reconstructed normalized expression with an *in silico* knocked-out gene, $p_t$, the perturbation effect for a cell type $t$ is given as $pef_t = \frac{p_t - b_t}{b_t}$. For this, we used Multicellular Transformer fine-tuned on the lung cancer atlas (Salcher et al. 2022) with normalized expression pseudobulked at cell type level. We have decided to use this model as this makes measuring the change in the normalized expression of a cell type easier.

###### A.3.7. Fine-tuning on lung cancer atlas

Multicellular Transformer can be fine-tuned allowing for dataset-specific analysis. We fine-tuned Multicellular Transformer on a lung cancer atlas (Salcher et al. 2022). The H5AD object was accessed from `https://datasets.cellxgene.cziscience.com/08adcdb7-1f28-42f6-8969-f8f13a2f82f5.h5ad` and split randomly by sample into train and validation following a 80%, 20% ratio. During fine-tuning, we leverage the pretrained gene embeddings and transformer block and train the gene expression embedding $MLP_e$, as well as gene expression reconstruction $MLP_r$ from scratch. This allows the model to adapt to the cell types present in the specific dataset, while reusing the information learned over the course of pretraining by leveraging pretrained gene identity embeddings, transformer block and masked gene classifier.

We have followed the same scRNA-seq normalization procedure as for pretraining corpus. We have set the nr of cells sampled from each sample to 512, therefore, the first layer of the $MLP_e$ used for fine-tuning has been also set to 512. Samples with less than 512 cells have been filtered out, and cell types which make up of less than 0.2% cell types were also filtered out. We have used the cell type labels present in the H5AD object downloaded from the CELLxGENE portal. The pretraining has been done using the same hyperparameters as during pretraining.

#### A.4. Datasets

##### A.4.1. Pretraining corpus

We have collected a corpus using 13 publicly available datasets from CELLxGENE (`https://cellxgene.cziscience.com/`). The dataset spanned more than 14M cells, including more than 3k donors and 4.2k samples. We have filtered out genes with less than 5 counts across cells and cells with less than 100 genes expressed using scanpy package (Wolf, Angerer, and Theis 2018). We have also filtered out samples with less than 1,024 cells, which is the number of cells sampled from each scRNA-seq sample. We took the union of all genes present across datasets, resulting in almost 60k genes and did not perform any highly variable genes selection, leveraging all genes. All of the data has been sequenced from *Homo Sapiens*. The corpus includes various tissues and diseases (Supp. Fig. S1) to allow for learning across diverse contexts. Exact list of datasets used and their statistics can be found in Supp. Table S1. We have harmonised the cell types across datasets and filtered out the cell types which made up less than 0.1% of all cells.

We have used the *CellArr* package (`https://github.com/BiocPy/cellarr`) to combine multiple datasets into one and allow seamless sampling of cells from the same sample during training and evaluation.

##### A.4.2. Lung cancer atlas

The lung cancer atlas (Salcher et al. 2022) has been accessed from the CELLxGENE portal (`https://datasets.cellxgene.cziscience.com/08adcdb7-1f28-42f6-8969-f8f13a2f82f5.h5ad`. We have filtered out genes and cells with less than 5 and 100 counts respectively using scanpy package. Additionally, we filtered out samples with less than 512 cells. We have used the cell type annotations provided in the dataset, filtering out cell types which made up less than 0.1% of all cells. We used the dataset for fine-tuning.

##### A.4.3. Spatial transcriptomics datasets

**Xenium human lung & breast cancer.** Both datasets have been downloaded from the 10x genomics website (lung cancer: `https://www.10xgenomics.com/datasets/ffpe-human-lung-cancer-data-with-human-immuno-oncology-profiling-panel-and-custom-add-on-1-standard`, breast cancer: https://www.10xgenomics.com/products/xenium-in-situ/preview-dataset-human-breast). We have filtered out genes and cells with less than 5 and 10 counts respectively using scanpy package.

For fine-tuning on the breast cancer dataset, we performed Leiden clustering (Traag, Waltman, and Van Eck 2019) using scanpy package (Wolf, Angerer, and Theis 2018) to annotate each cell with a label. We have then used these cluster labels as the equivalent of cell type labels during fine-tuning. Therefore, showing how one can use alternative labels to cell type labels for training the model.

**Visium glioblastoma**. The Visium glioblastoma dataset has been accessed using scanpy package (sample id: `Parent_Visium_Human_Glioblastoma`). We have filtered out genes with less than 10 counts across cells and cells with less than 1000 or more than 35000 counts respectively using scanpy.

###### A.4.4. DepMap

We have extracted the publicly available viability scores for each gene from the DepMap consortium (Tsherniak et al. 2017) website (https://depmap.org/portal/data_page/). We leveraged the `CRISPRGeneEffect.csv` file to extract gene effect estimates for all cell lines. The gene effect estimates have been already copy number corrected, scaled, and screen quality corrected. We filtered to relevant lung cancer cell lines present in both DepMap as well as lung cancer atlas, specifically *lung adenocarcinoma, squamous cell lung carcinoma and non-small cell lung carcinoma*, leaving us with gene effect estimates for 81 cell lines across 18443 genes.

#### Appendix B. Supplementary Figures & Tables

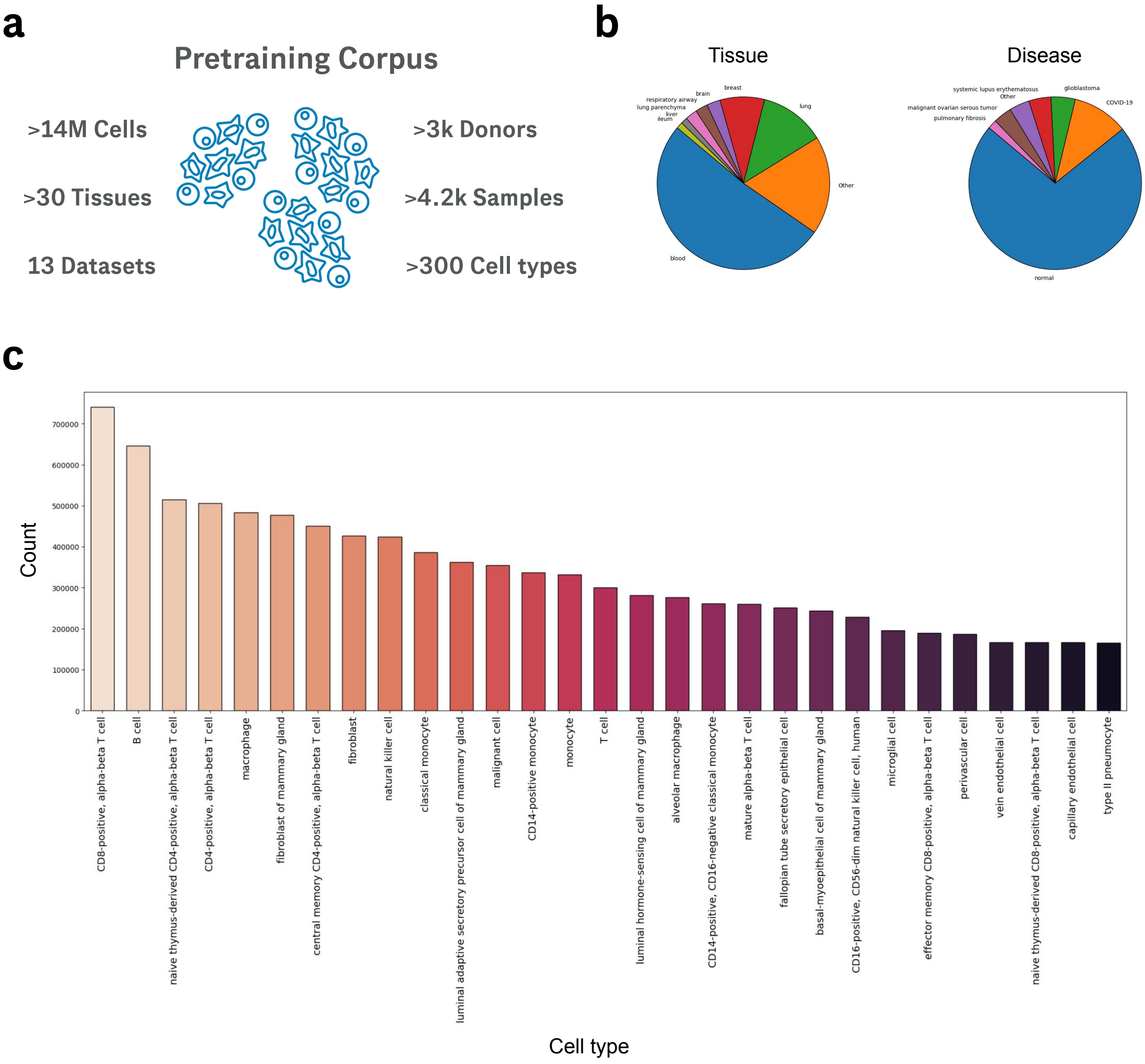

FIGURE S1. **Pretraining corpus statistics (a)** Overall statistics of the pretraining corpus. **(b)** Distribution of tissues and diseases in the pretraining corpus. **(c)** Most common cell type counts present in the pretraining corpus across datasets.

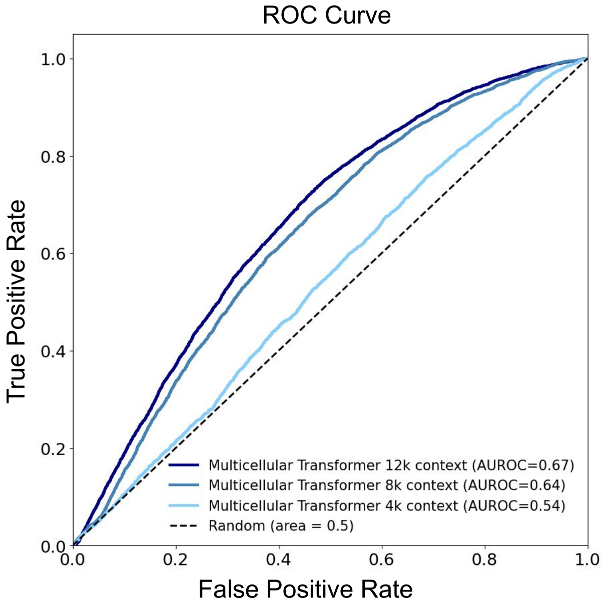

FIGURE S2. **Impact of the context size on the performance of the Multicellar Transformer.** Ablation study showing the comparison of Multicellular Transformer model pretrained with various context size (i.e. number of genes) on a dataset of known receptor-ligand interactions and randomly sampled negatives (Methods).

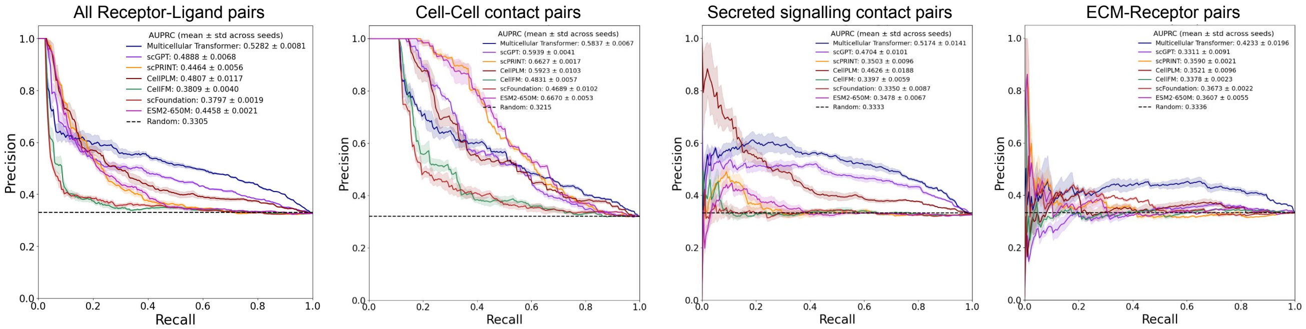

FIGURE S3. **Additional results on the zero-shot receptor-ligand identification task.** Performance on the receptor-ligand prediction task using pretrained gene embeddings (zero-shot) across methods as measured by an area under the precision-recall curve (AUPRC). The true receptor-ligand pairs were taken from CellChatDB receptor-ligand set Jin et al. (2021), while the negatives were randomly sampled with different seeds (Methods). Scores for each receptor-ligand pair were computed by calculating cosine similarity between embeddings coming from each model. We stratify the results by the interaction type using the annotations from CellChatDB.

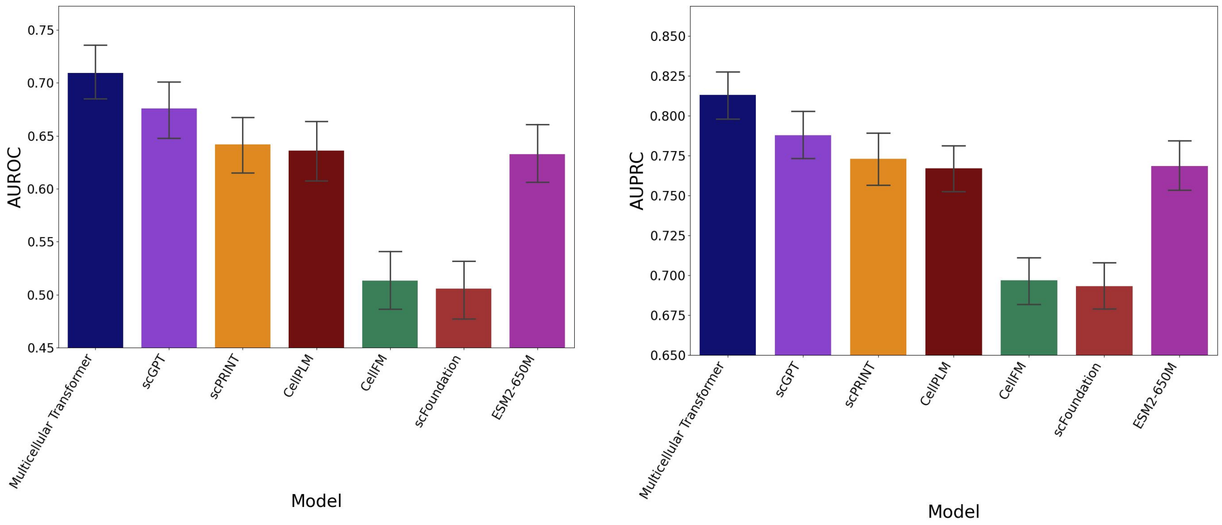

FIGURE S4. **Zero-shot performance across receptors on the receptor-ligand identification task.** Performance on the receptor-ligand prediction task across receptors using pretrained gene embeddings (zero-shot) across methods as measured by an area under the receiver operating characteristic (AUROC; left) and precision-recall curves (AUPRC; right). The true receptor-ligand pairs were taken from CellChatDB receptor-ligand set Jin et al. (2021), while the negatives were randomly sampled with different seeds (Methods). Scores for each receptor-ligand pair were computed by calculating cosine similarity between embeddings coming from each model. The metrics were then computed for each receptor. The error bars represent 95% confidence interval.

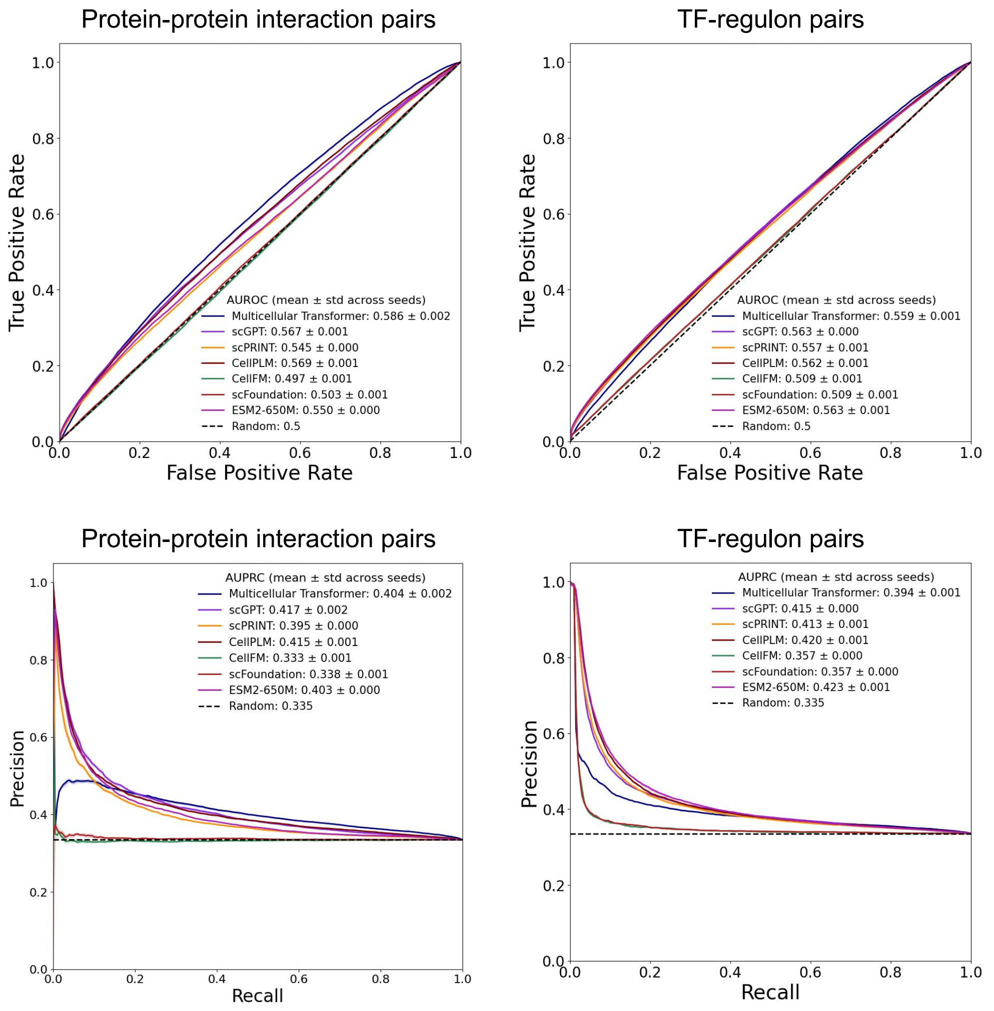

FIGURE S5. **PPI & TF-regulon zero-shot identification task results.** Performance on the PPI (left) and TF-regulon (right) prediction task using pretrained gene embeddings (zero-shot) across methods as measured by an area under the receiver operating characteristic (AUROC; top) and precision-recall curves (AUPRC; bottom). The true interaction pairs were taken from OmniPath (Türei, Korcsmáros, and Saez-Rodriguez 2016) (PPI) and CollecTRI (Müller-Dott et al. 2023) (TF-regulon), while the negatives were randomly sampled with different seeds (Methods). Scores for each pair were computed by calculating cosine similarity between embeddings coming from each model.

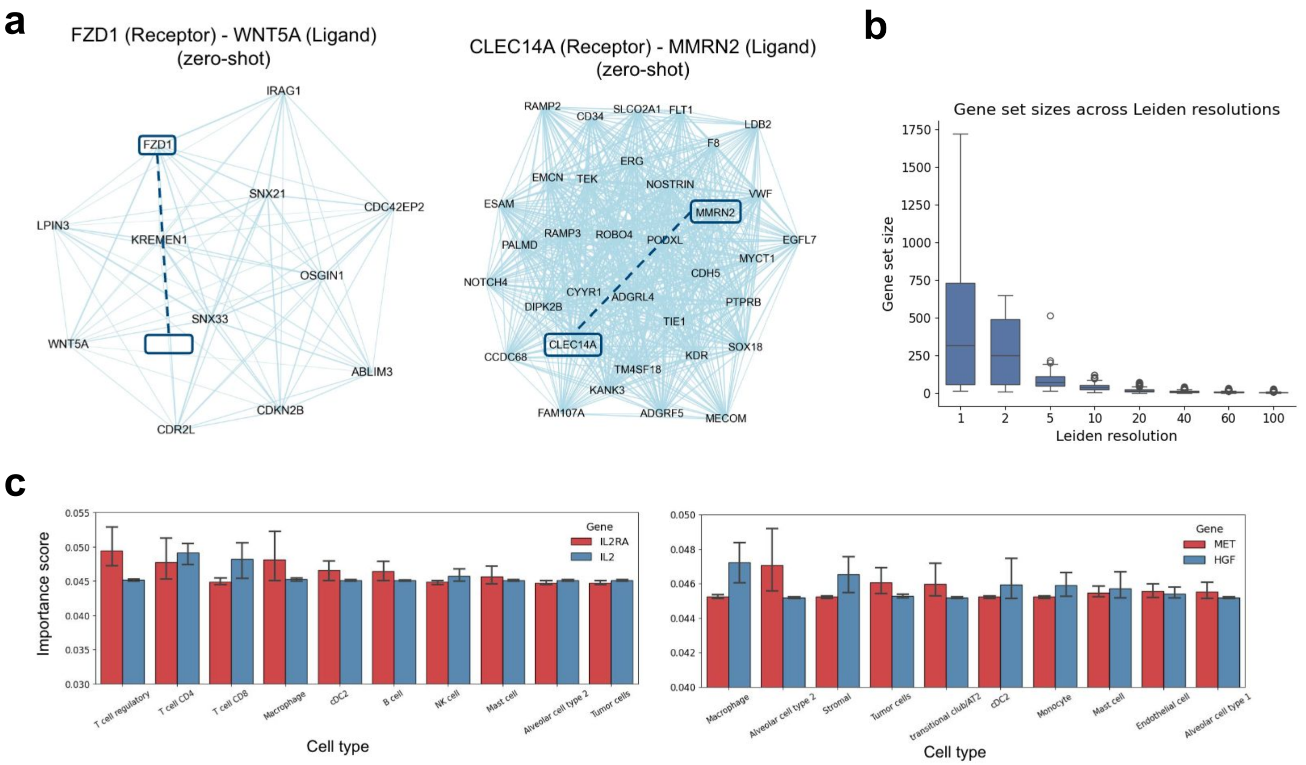

FIGURE S6. **Analysis of the zero-shot gene sets with receptor-ligand pairs. (a)** Examples of the zero-shot gene sets identified by the Multicellular Transformer model at the Leiden resolution of 50 with receptor-ligand pairs. **(b)** Most important cell types for *IL2RA-IL2R* and *MET-HGF* receptor-ligand pairs using integrated gradients (Sundararajan, Taly, and Yan 2017) across validation samples on lung cancer atlas (Salcher et al. 2022). Cell type importance scores have been computed by applying softmax function to aggregated absolute attribution scores across samples. The error bars represent 95% confidence interval. cDC stands for conventional dendritic cells.

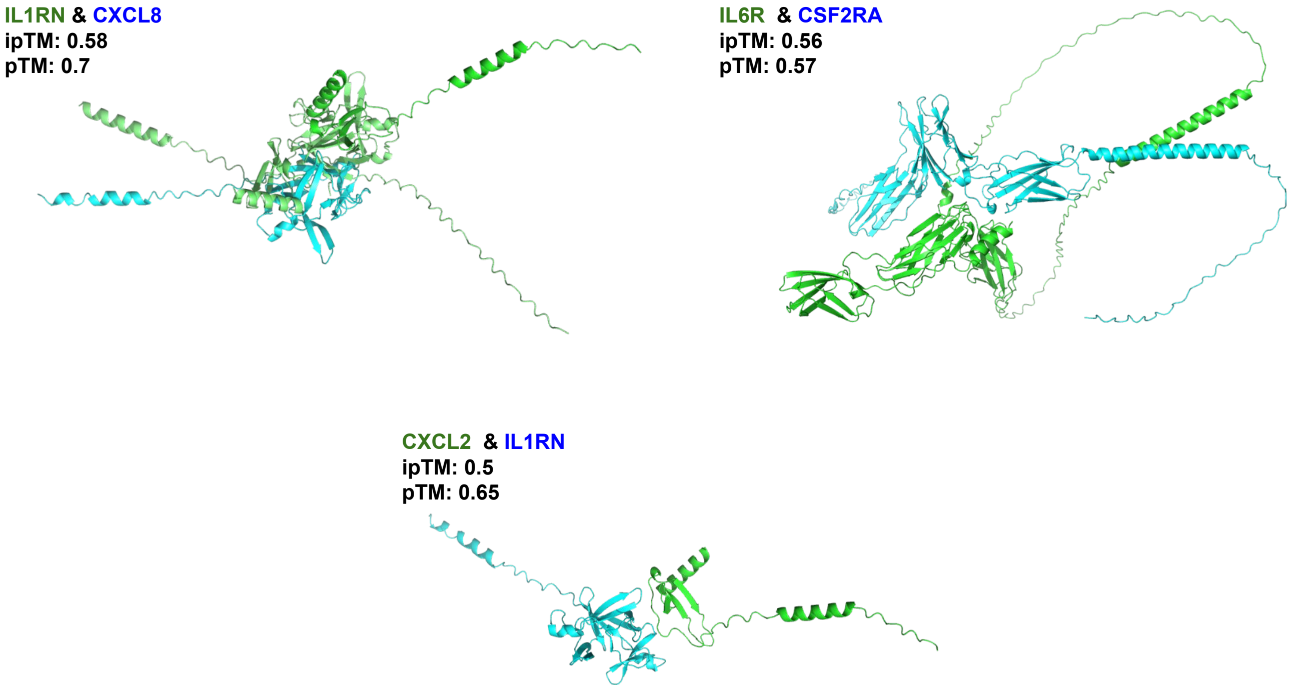

FIGURE S7. **Predicted protein complex structures identified with Multicellular Transformer** Structures of protein complexes predicted with AlphaFold3 (Abramson et al. 2024) and deemed to interact with Multicellular Transformer. The pTM (predicted Template Modeling) score measure how well different parts of protein fold together, while ipTM (inter-chain pTM) is the adaptation of pTM for assessing the structural confidence of protein-protein interfaces, higher is better.

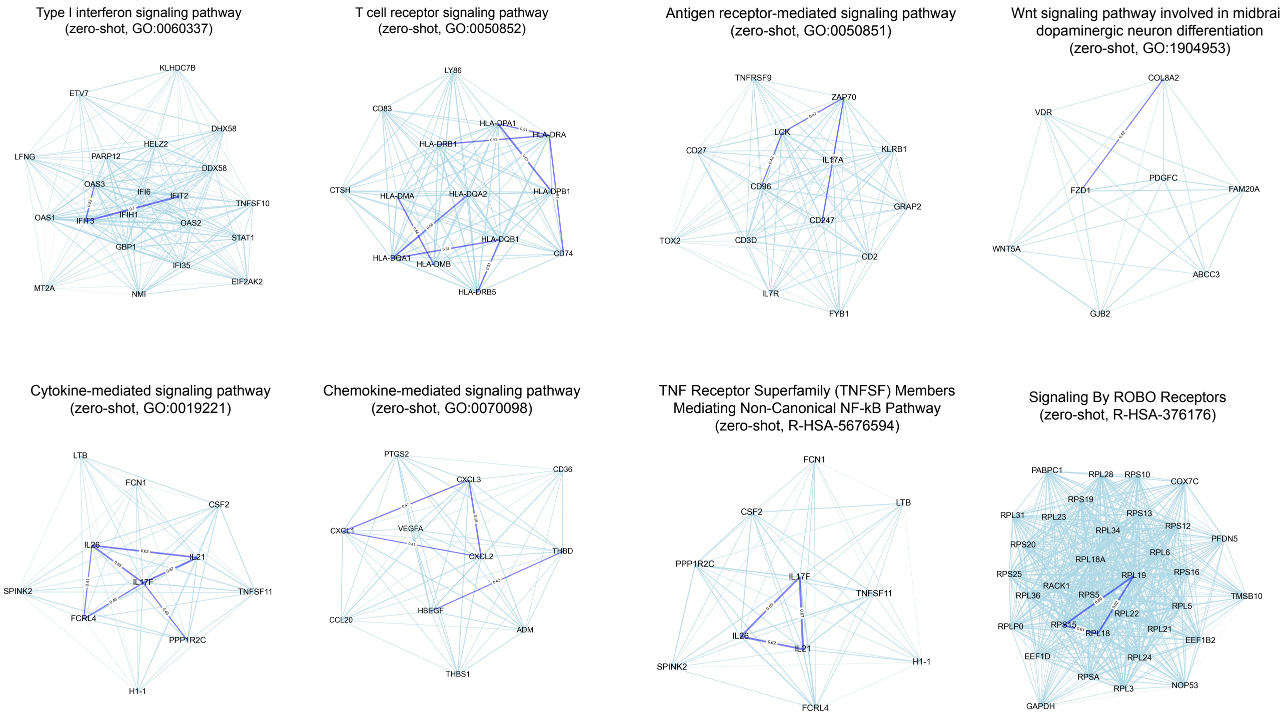

FIGURE S8. **Zero-shot gene sets identified with Multicellular Transformer** Examples of the zero-shot gene sets identified by the Multicellular Transformer model at the Leiden resolution of 50 together with pathway enriched in the corresponding gene set.

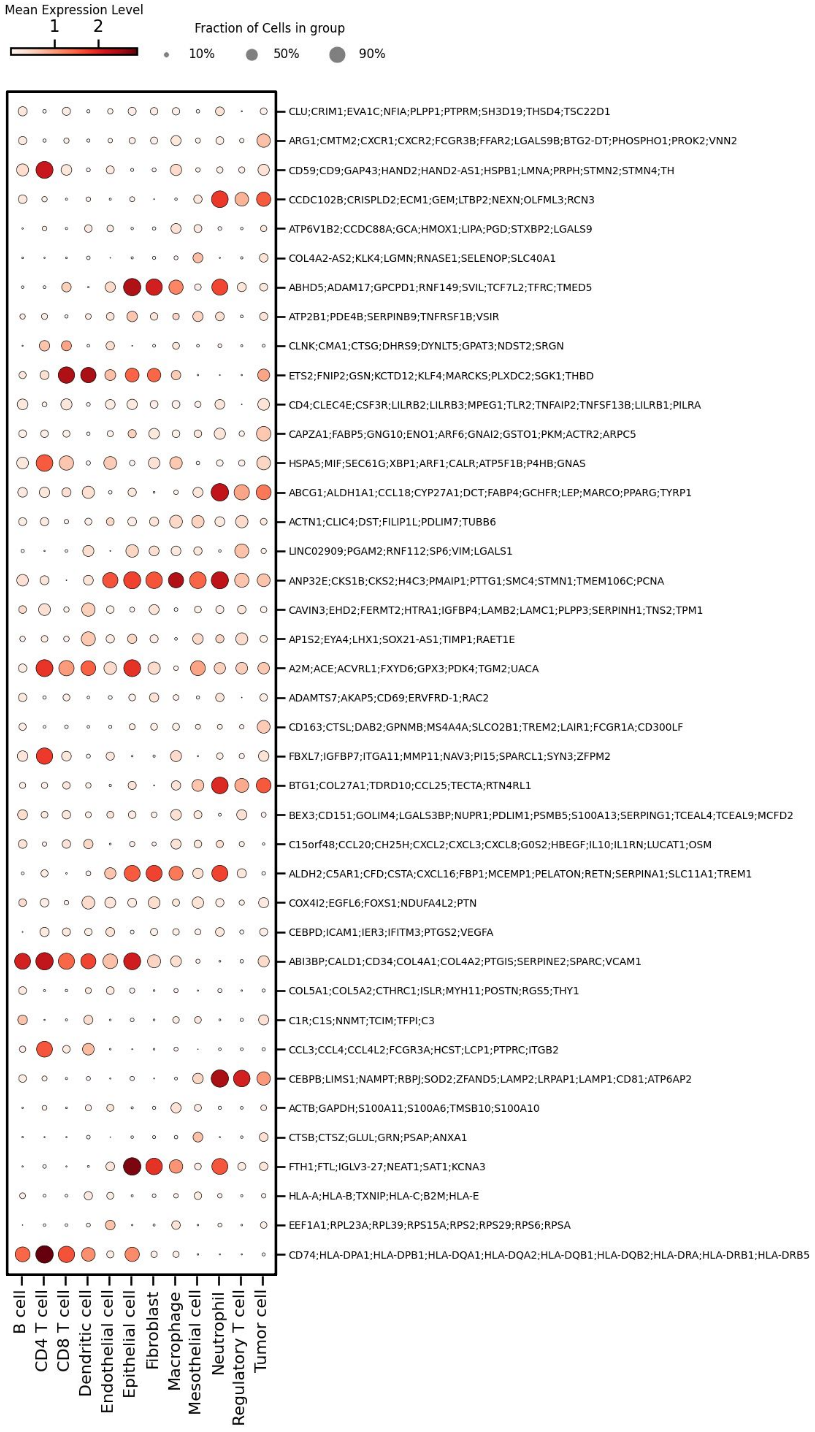

FIGURE S9. **Gene sets identified with Multicellular Transformer.** Normalized expression of a set of gene sets identified by the Multicellular Transformer at Leiden resolution of 50 across a set of cell types in lung cancer atlas (Salcher et al. 2022).

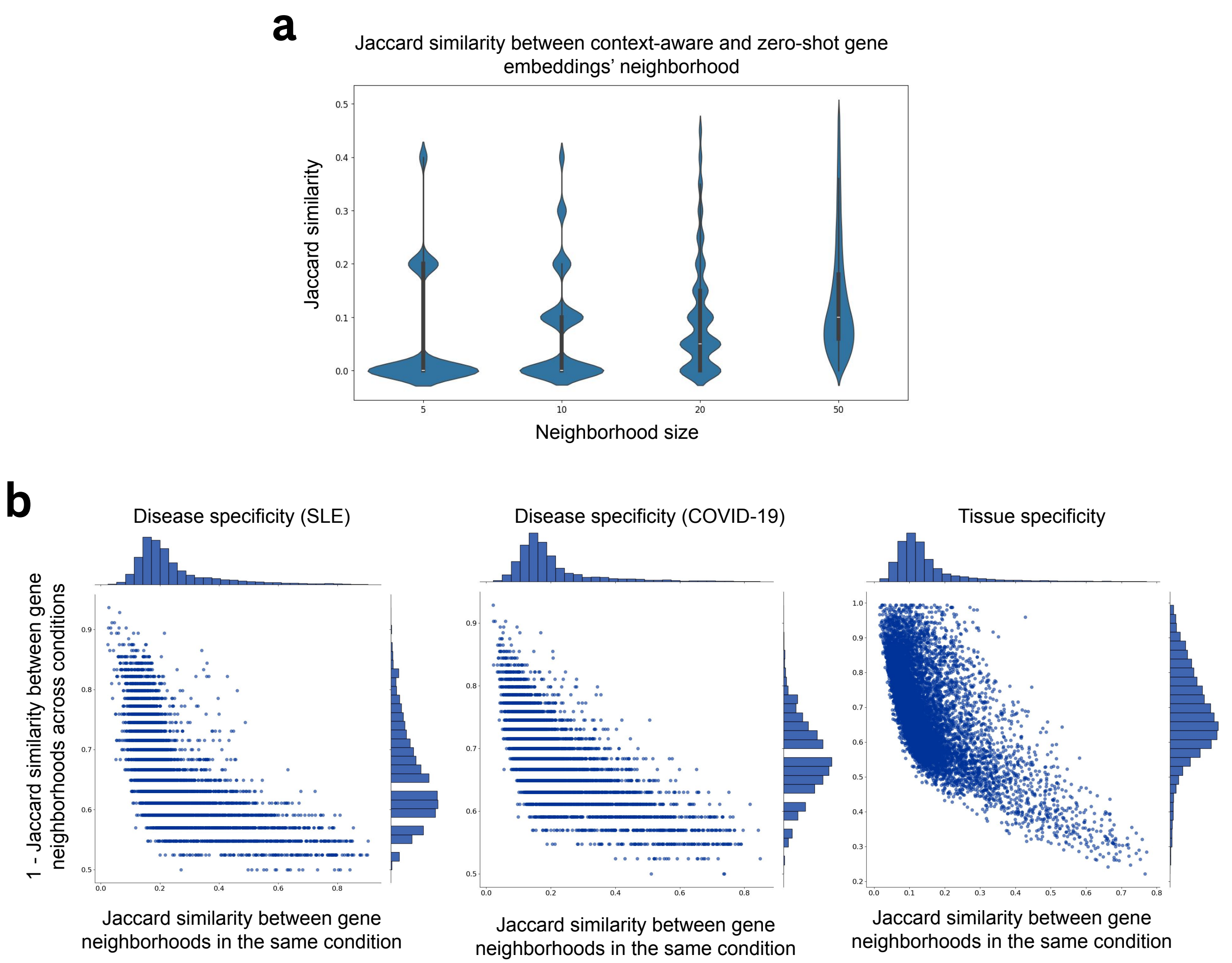

FIGURE S10. **Multicellular Transformers identifies condition-specific genes using gene neighborhood across samples. (a)** The Jaccard similarity between context-aware and zero-shot gene embeddings' neighborhood at different sizes of the neighborhood across pretraining samples. **(b)** Pairwise plots showing the mean overlap between gene neighborhoods across healthy vs disease samples in SLE, COVID-19 and tissues. The overlap was calculated by using 20 nearest neighbors for each gene in each sample as measured by cosine similarity between gene embeddings.

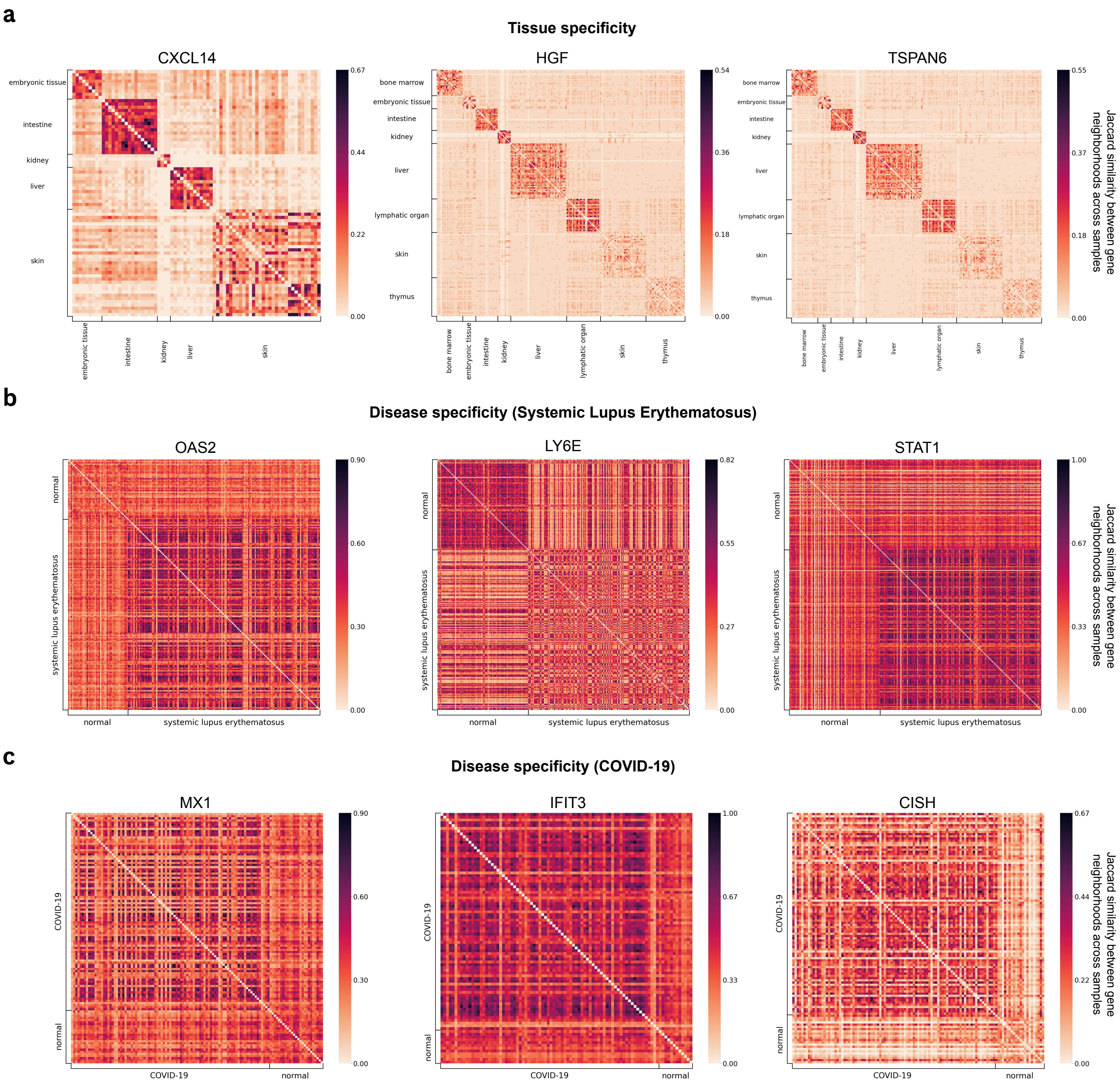

FIGURE S11. **Additional analysis of context-aware gene embeddings (a)** Genes with high condition-specificity across tissues and diseases (Lupus & COVID-19). The condition-specificity was calculated by dividing the mean overlap between gene neighborhoods in the same condition by the mean overlap between gene neighborhoods across conditions. The datasets used to calculate condition specificity were Suo et al. (2022) for tissue analysis, Perez et al. (2022) for lupus and Stephenson et al. (2021) for COVID-19.

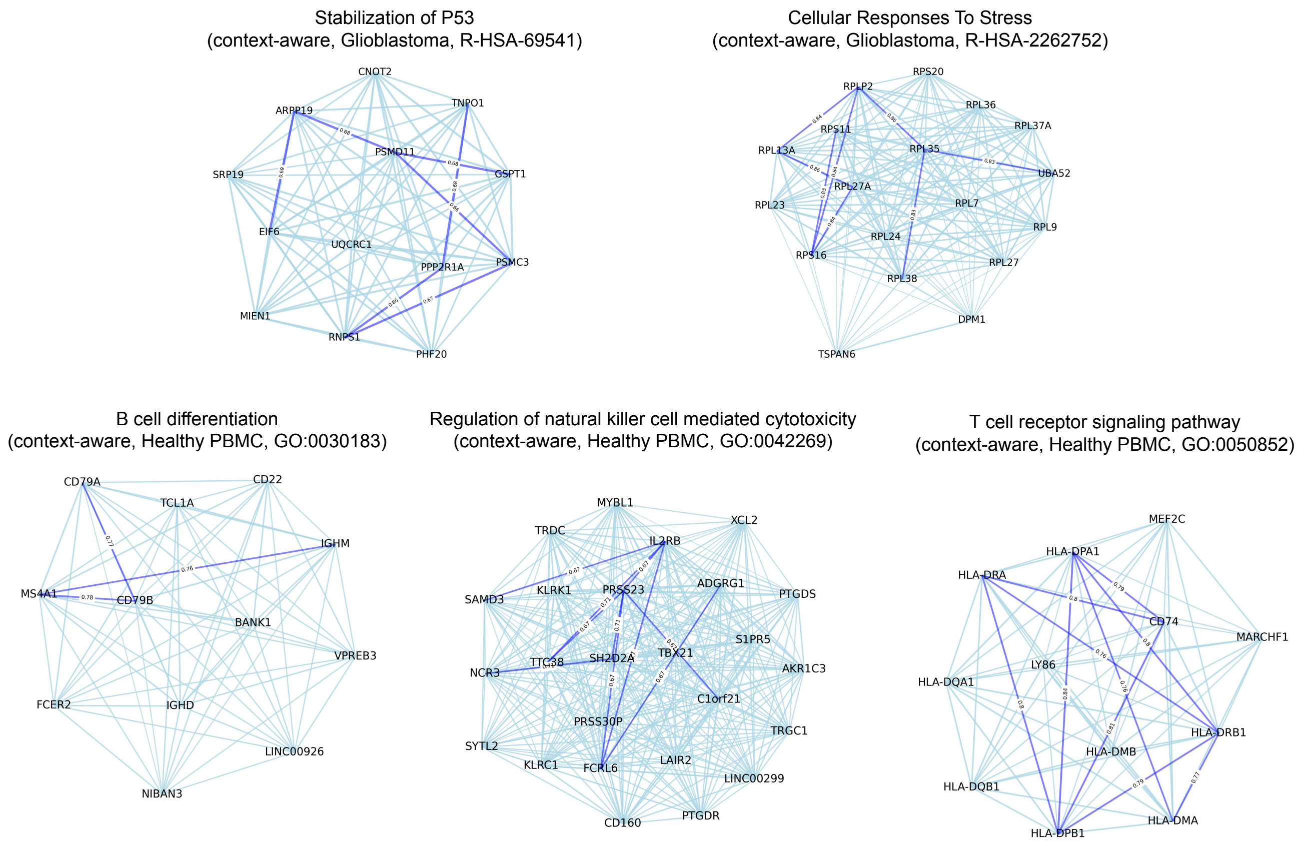

FIGURE S12. **Context-aware gene sets identified with Multicellular Transformer gene embeddings.** Examples of the context-aware gene sets identified by the Multicellular Transformer model at the Leiden resolution of 50 together with pathway enriched in the corresponding gene set in Glioblastoma (top) and healthy PBMC (bottom) samples. The samples were extracted from Glioblastoma atlas (Ruiz-Moreno et al. 2022) and single-cell eQTL mapping study (Yazar et al. 2022) respectively.

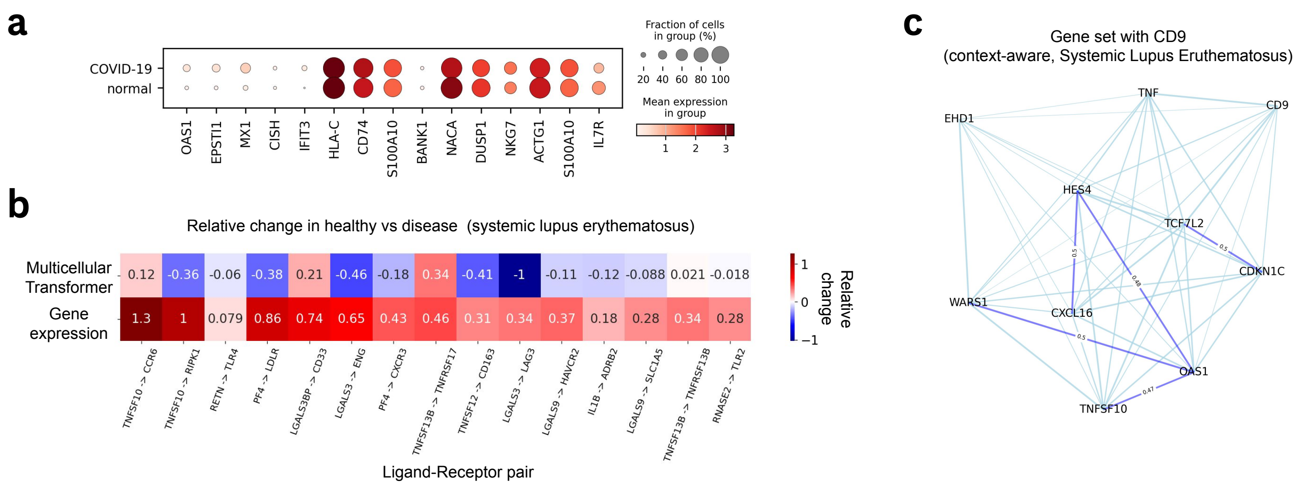

FIGURE S13. **Expression of disease-specific genes in COVID-10 identified with Multicellular Transformer & receptor-ligand analysis in systemic lupus erythematosus. (a)** The expression of disease-specific genes in COVID-19 (Stephenson et al. 2021) computed using context-aware embeddings from Multicellular Transformer. Many genes have been shown to be associated with the disease but do not exhibit a significant change in expression between healthy and disease. **(b)** Relative change in cosine similarity (top) vs expression (bottom) in healthy vs disease patients with SLE across a set of known receptor-ligand pairs with highest change according to co-expression in (Perez et al. 2022). **(c)** Context-aware gene set in SLE (Perez et al. 2022) with CD9 gene. The gene set has been discovered by clustering context-aware gene embeddings from SLE samples at Leiden resolution of 50.

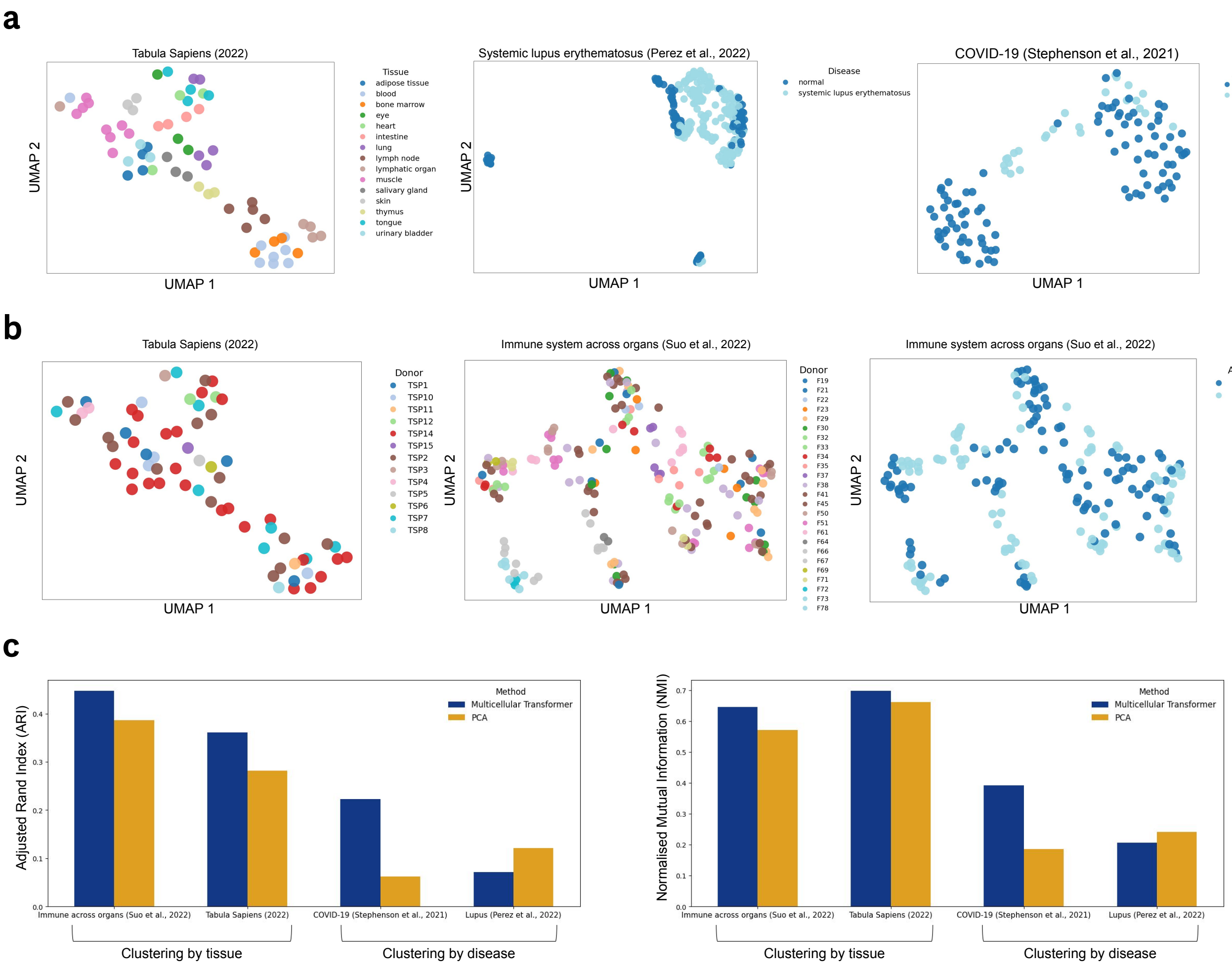

FIGURE S14. **Further results on sample-level clustering (a)** UMAPs of sample-level embeddings across tissues on the Tabula Sapiens (Consortium* et al. 2022) dataset (left) and SLE (Perez et al. 2022) (middle) and COVID-19 (Stephenson et al. 2021). The sample embeddings were extracted by averaging out all of the contextual gene embeddings in the sample. **(b)** UMAPs of sample-level embeddings across donors on the Tabula Sapiens (left) (Consortium* et al. 2022) and across donors (middle) and assays (right) on immune across organs dataset (Suo et al. 2022). The sample embeddings were extracted by averaging out all of the contextual gene embeddings in the sample. **(c)** Performance across clustering metrics of Multicellular Transformer vs PCA baseline. For both Adjusted Rand Index (ARI) and normalized Mutual Information (NMI) higher is better, minimum is 0 and maximum is 1.

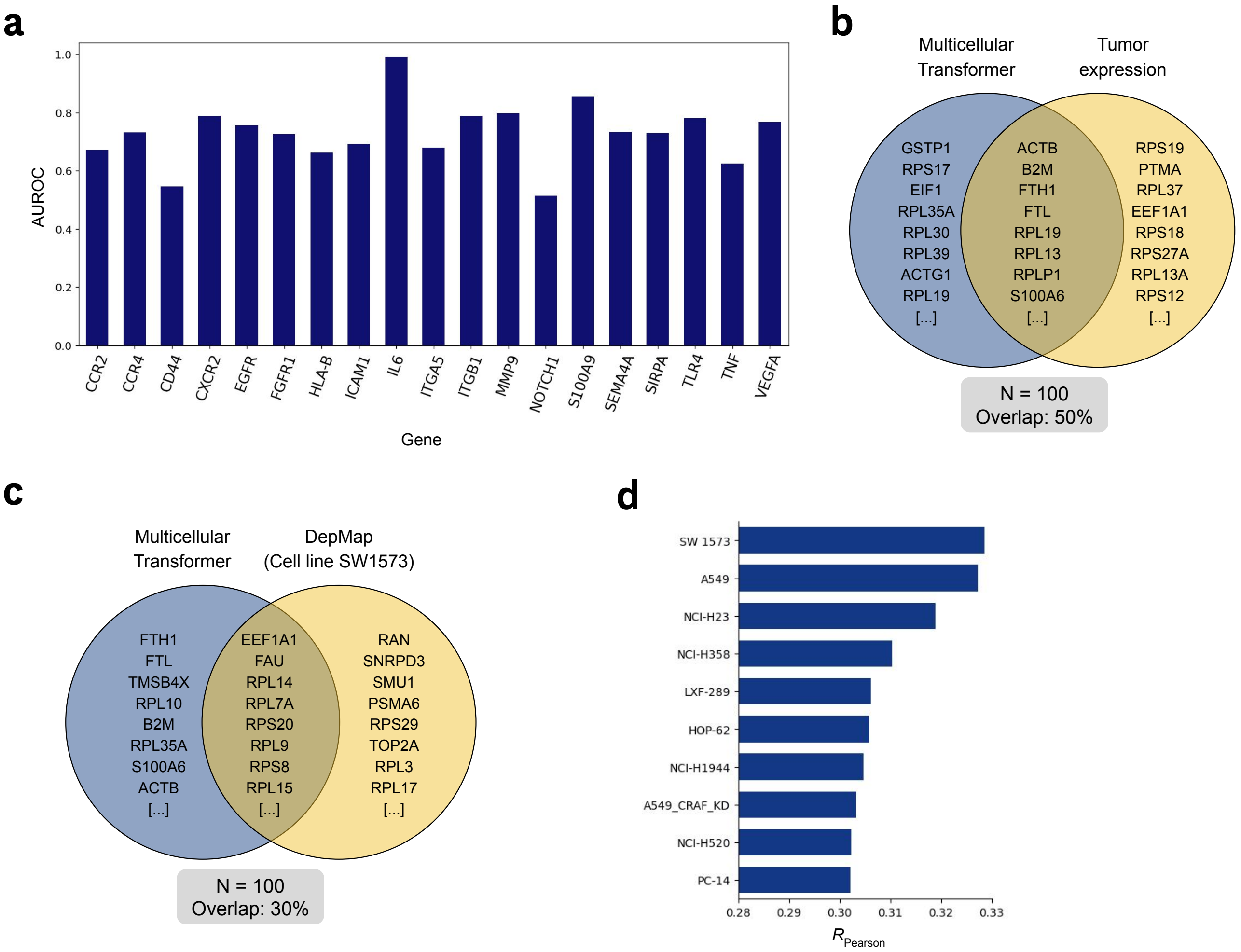

FIGURE S15. ***In silico* genetic perturbation effect estimation at cell population level.** **(a)** AUROC per gene on the *in silico* gene knock-out effect on a set of true interacting genes vs control genes. The control genes were randomly selected from a pool of available genes. The knock-out effect is quantified by the maximum log-fold change in gene's logits (Methods). **(b)** The overlap between genes with highest effect from Multicellular Transformer (i.e. one which leads to highest reduction in the normalized tumor expression) and genes with highest normalized tumor expression. **(c)** The overlap between genes with highest effect from Multicellular Transformer (i.e. one which leads to highest reduction in the normalized tumor expression) and from viability readouts following CRISPR knock-outs from DepMap (Tsherniak et al. 2017) data across lung cancer cell lines. **(d)** The lung cancer cell lines with highest pearson correlation between the viability readouts following CRISPR knock-outs from DepMap (Tsherniak et al. 2017) and the predicted normalized effect from Multicellular Transformer.

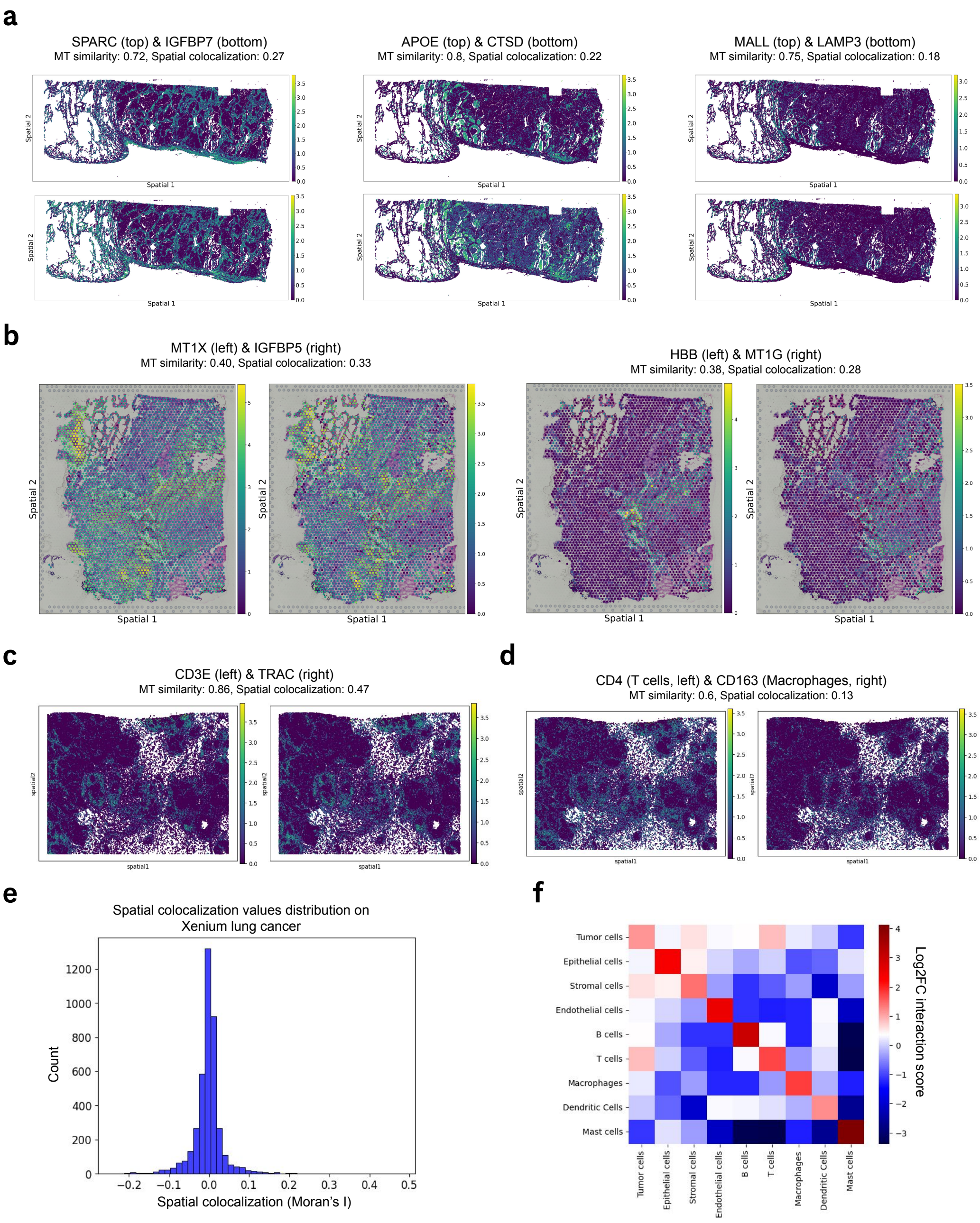

FIGURE S16. **Additional results on spatial colocalization patterns across datasets recovered by the Multicellular Transformer (a)** Spatial colocalization of selected gene pairs in Xenium human lung cancer dataset, recovered by the Multicellular Transformer in a zero-shot fashion by assessing genes with highest cosine similarity in gene embeddings for each gene. The spatial colocalization was measured by computing bivariate Moran's I between two genes. **(b)** Spatial colocalization of selected gene pairs in Visium glioblastoma dataset, recovered by the Multicellular Transformer in a zero-shot fashion by assessing genes with highest cosine similarity in gene embeddings for each gene. The spatial colocalization was measured by computing bivariate Moran's I between two genes. **(c)** Spatial colocalization of *CD3E* & *TRAC* genes in Xenium breast cancer dataset, recovered by the Multicellular Transformer fine-tuned on the dataset by assessing genes with highest cosine similarity in the gene embedding space. **(d)** Spatial colocalization of *CD4* & *CD163* genes in Xenium breast cancer dataset, recovered by the Multicellular Transformer fine-tuned on the dataset by assessing T cell-Macrophage gene pairs with highest cosine similarity in the gene embedding space. **(e)** The distribution of spatial colocalization values as measured by bivariate Moran's I on Xenium lung cancer dataset. **(f)** Interaction matrix between cell types computed using the gene co-expression between marker genes on the Xenium human breast cancer dataset. The interaction score was computed by calculating the *log*2 fold change between the gene pair's co-expression divided by its average co-expression across all genes and taking the average across gene pairs for each cell type-cell type pair.

| Study | # Cells | # Samples |
|---|---|---|
| AIDA Phase 1 Data Freeze v1: Chinese, Indian, Japanese, Korean, and Malay donors in Japan, Singapore, and South Korea (Kock et al. 2024) | 1,058,909 | 508 |
| COVID-19 immune features revealed by a large-scale single-cell transcriptome atlas (Ren et al. 2021) | 1,462,702 | 284 |
| Single-cell multi-omics analysis of the immune response in COVID-19 (Stephenson et al. 2021) | 647,366 | 143 |
| Harmonized single-cell landscape, intercellular crosstalk and tumor architecture of glioblastoma (Ruiz-Moreno et al. 2022) | 1,060,257 | 191 |
| Human breast cell atlas (Reed et al. 2024) | 2,122,065 | 351 |
| Cells of the adult human heart (Litviňuková et al. 2020) | 486,134 | 88 |
| Cells of the human intestinal tract mapped across space and time (Elmentaite et al. 2021) | 428,469 | 159 |
| The integrated Human Lung Cell Atlas (Sikkema et al. 2023) | 2,282,120 | 731 |
| Mapping the developing human immune system across organs (Suo et al. 2022) | 908,046 | 221 |
| Single-cell RNA-seq reveals the cell-type-specific molecular and genetic associations to lupus (Perez et al. 2022) | 1,263,676 | 274 |
| MSK SPECTRUM – Ovarian cancer mutational processes drive site-specific immune evasion (Vázquez-García et al. 2022) | 929,690 | 146 |
| Single-cell eQTL mapping identifies cell type specific genetic control of autoimmune disease (Yazar et al. 2022) | 1,248,980 | 981 |
| Tabula Sapiens (Consortium* et al. 2022) | 483,152 | 129 |
| **Total** | **14,381,566** | **4,206** |

TABLE S1. Summary of studies used in pretraining corpus and their respective cell and sample counts.

| Cell Type | Marker Genes |
|---|---|
| Tumor cells | ABCC11, AGR3, DSP, ERBB2, FOXA1, GATA3, S100A14, SERPINA3, TACSTD2, AGR3, CEACAM6, MZB1, SERPINA3, FASN |
| Epithelial cells | ACTA2, DST, KRT14, KRT5, MYLK, KRT15, KRT23, PTN, SFRP1 |
| Stromal cells | ALDH1A3, GJB2, LUM, POSTN, SFRP4 |
| Endothelial cells | AQP1, EGFL7, PECAM1, VWF |
| B cells | BANK1, CD79A, MS4A1 |
| T cells | CCL5, CD4, CD8A, CXCR4, CYTIP, IL7R, LTB, TRAC |
| Macrophages | APOC1, C15orf48, C1QA, C1QC, CD14, CD163, CD68, FGL2, ITGAX, MMP12 |
| Dendritic Cells | CCR7, CD83, IL3RA, LILRA4, PLD4 |
| Mast cells | CPA3, CTSG, KIT |

TABLE S2. Marker genes for each cell type used in Xenium breast cancer analysis.

### Appendix C. Sensitivity of contextual embeddings to cell ordering and resampling

To assess whether MCT contextual gene embeddings are sensitive to the ordering or identity of cells assigned to the expression encoder, we performed an input perturbation analysis using three lung cancer samples. Starting from an unperturbed reference input, we perturbed increasing fractions of cell positions (0–100%) using three schemes: **(i) within-cell-type shuffling**, in which cell positions were permuted only within their respective cell-type blocks; **(ii) cell resampling**, in which cells were replaced by newly sampled cells from the same donor and cell type; and **(iii) across-cell-type shuffling**, in which cells were permuted across cell-type blocks, thereby violating the expected cell-type slot assignments. For each perturbation, we quantified contextual embedding drift as the mean cosine distance from the corresponding embeddings obtained from the unperturbed input.

Within-cell-type permutation produced virtually no change in the contextual embeddings, with mean cosine distance remaining below 0.005 even when all cell positions were shuffled (Fig. S17). Resampling cells from the same donor and cell type produced a very modest drift, reaching approximately 0.035 at complete replacement. In contrast, shuffling cells across cell-type blocks produced substantially greater and progressively increasing drift, reaching approximately 0.134 when all positions were perturbed. For context, these changes remain considerably smaller than the mean cosine distance between a gene and its nearest neighboring gene in the embedding space (0.472) or between randomly selected gene pairs (0.834).

These results demonstrate that contextual embeddings are robust to arbitrary cell ordering *within* a cell type and resampling, while remaining sensitive to the cell-type identity associated with each input block. This behavior is consistent with the pretraining procedure, in which cells are randomly sampled at each training epoch, explicitly exposing the model to different cells and within-cell-type arrangements throughout training.

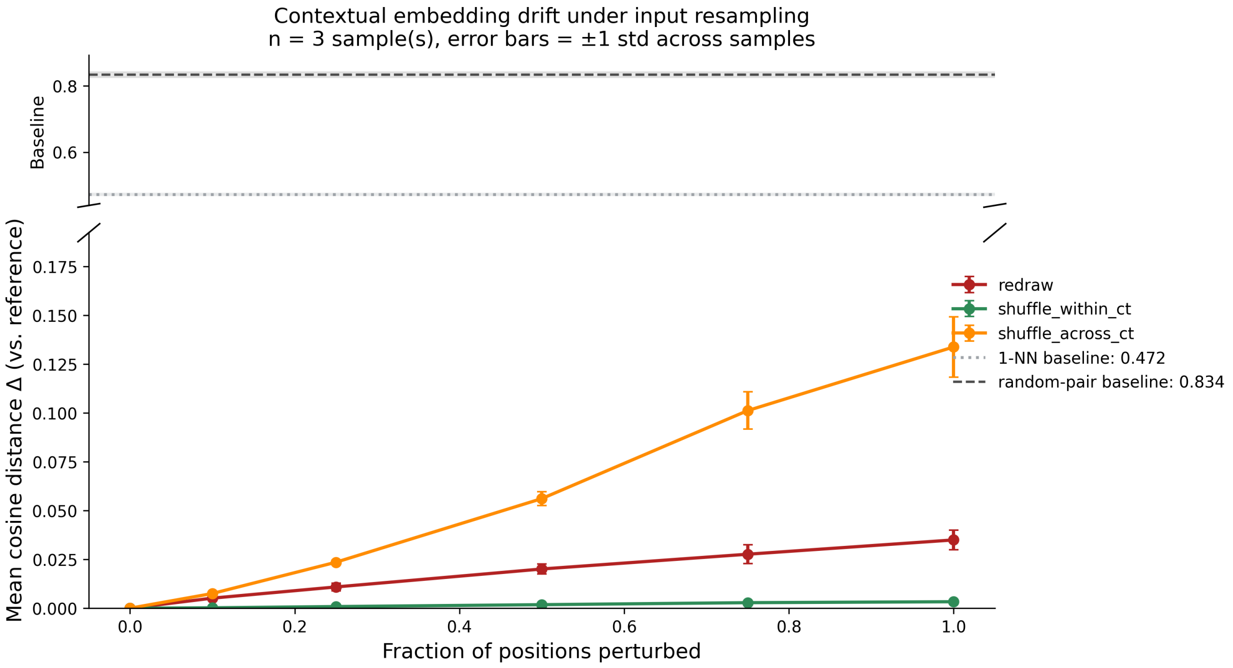

FIGURE S17. **Sensitivity of contextual gene embeddings to cell ordering and resampling.** Contextual embedding drift was measured as the mean cosine distance relative to embeddings obtained from the unperturbed reference input across three lung cancer samples (mean ± 1 standard deviation). Increasing fractions of cell positions were perturbed by shuffling cells within their respective cell-type blocks (`shuffle_within_ct`), resampling cells from the same donor and cell type (`redraw`), or shuffling cells across cell-type blocks (`shuffle_across_ct`). Within-cell-type shuffling results in negligible embedding drift even when all positions are perturbed, whereas resampling cells produces only modest drift. In contrast, shuffling cells across cell-type blocks produces substantially greater drift, indicating sensitivity to the cell-type identity assigned to each input block. Dashed horizontal lines show the mean cosine distance between a gene and its nearest neighboring gene (1-NN; 0.472) and between random gene pairs (0.834) in the contextual embedding space. The broken $y$-axis is used to display these embedding-space reference distances alongside the substantially smaller perturbation-induced changes.

### Appendix D. Scaling Laws

We evaluated how MCT performance scales with the number of independent biological samples available during pretraining. We trained MCT on four subsets corresponding to 5%, 10%, 25%, and 50% of the available pretraining data, comprising 47, 94, 236, and 472 biological samples, respectively. Model architecture, optimizer, masking strategy, and validation and test sets were held fixed across all runs.

To isolate the effect of increasing the amount of independent pretraining data, we kept the number of exposures per sample fixed rather than the total number of optimization steps. Specifically, each model was trained for two epochs, such that every biological sample was observed twice. Consequently, models trained on larger datasets receive proportionally more optimization steps because they are exposed to more unique samples. Holding the number of optimization steps fixed instead would cause samples in the smaller datasets to be seen more frequently, confounding the effect of dataset size with repeated exposure to the same biological samples.

We use validation loss as the measure of model performance. Prior work on neural scaling laws, particularly in the language-modeling domain, has established held-out loss as a useful measure for characterizing how model performance changes with training data and compute (Kaplan et al. 2020). Following this framework, we examined the relationship between the number of pretraining samples and validation loss using power-law fits of the form $L(D) = AD^{-\alpha}$, where $D$ denotes the number of independent biological samples.

Increasing the amount of pretraining data improved validation performance across the different training objectives (Fig. S18). The power-law fits explained a substantial fraction of the variation across dataset sizes ($R^2 = 0.66$–$0.83$), although the estimated scaling exponents were relatively small ($\alpha = 0.012$–$0.051$).

Notably, the largest improvements occurred as the dataset increased from 5% to 25% of the available samples, while increasing the dataset from 25% to 50% produced little additional improvement (Fig. S19). These results indicate that MCT benefits from increasing the number of independent biological samples during pretraining, but exhibits diminishing returns at the upper end of the range examined.

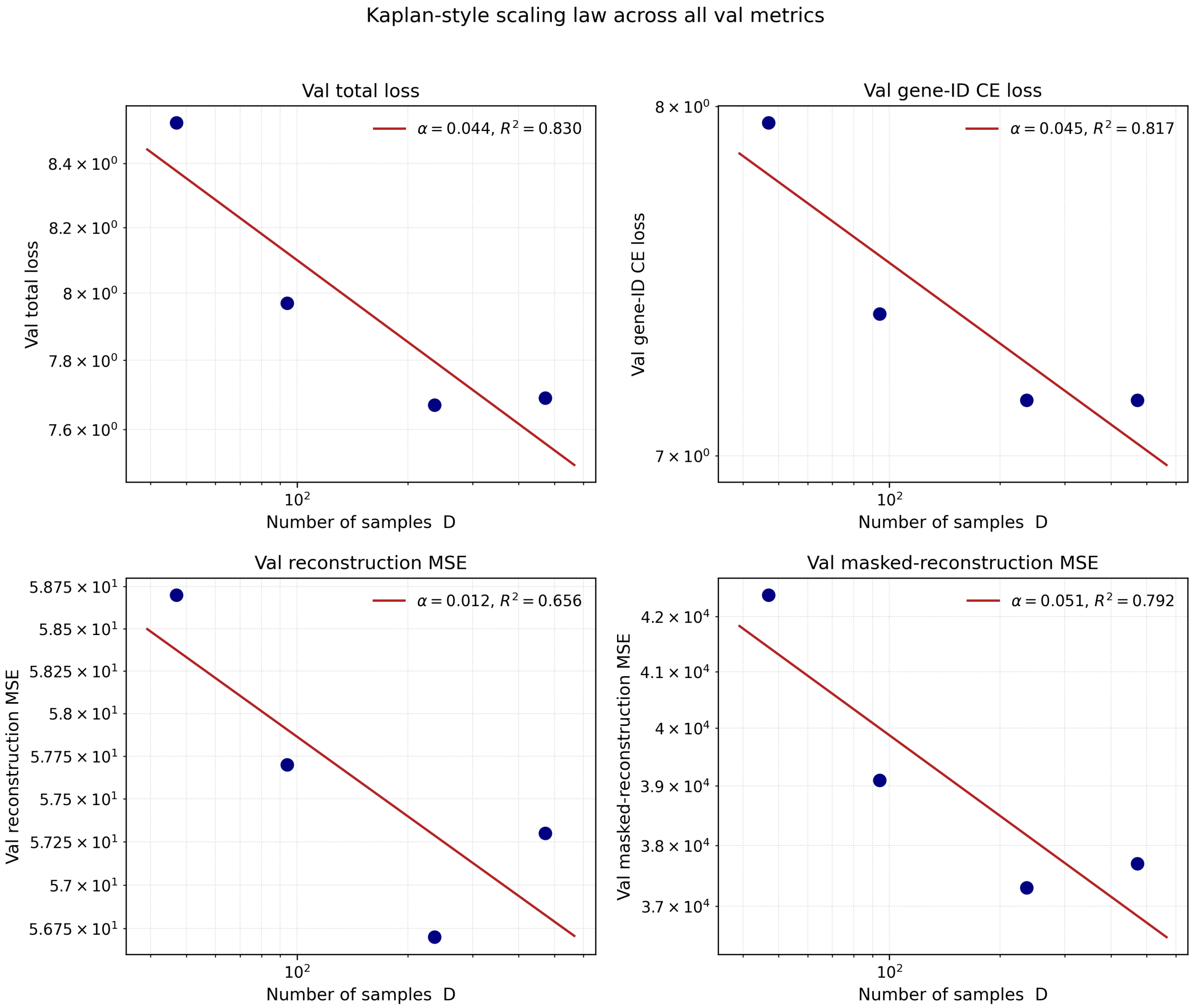

FIGURE S18. **Scaling of MCT validation performance with the number of pretraining samples.** Validation total loss, gene-ID cross-entropy loss, reconstruction MSE, and masked-reconstruction MSE for models pretrained using 47, 94, 236, and 472 independent biological samples. Red lines show power-law fits of the form $L(D) = AD^{-\alpha}$. All models use the same architecture and training configuration and are trained for two epochs, ensuring equal exposure per biological sample.

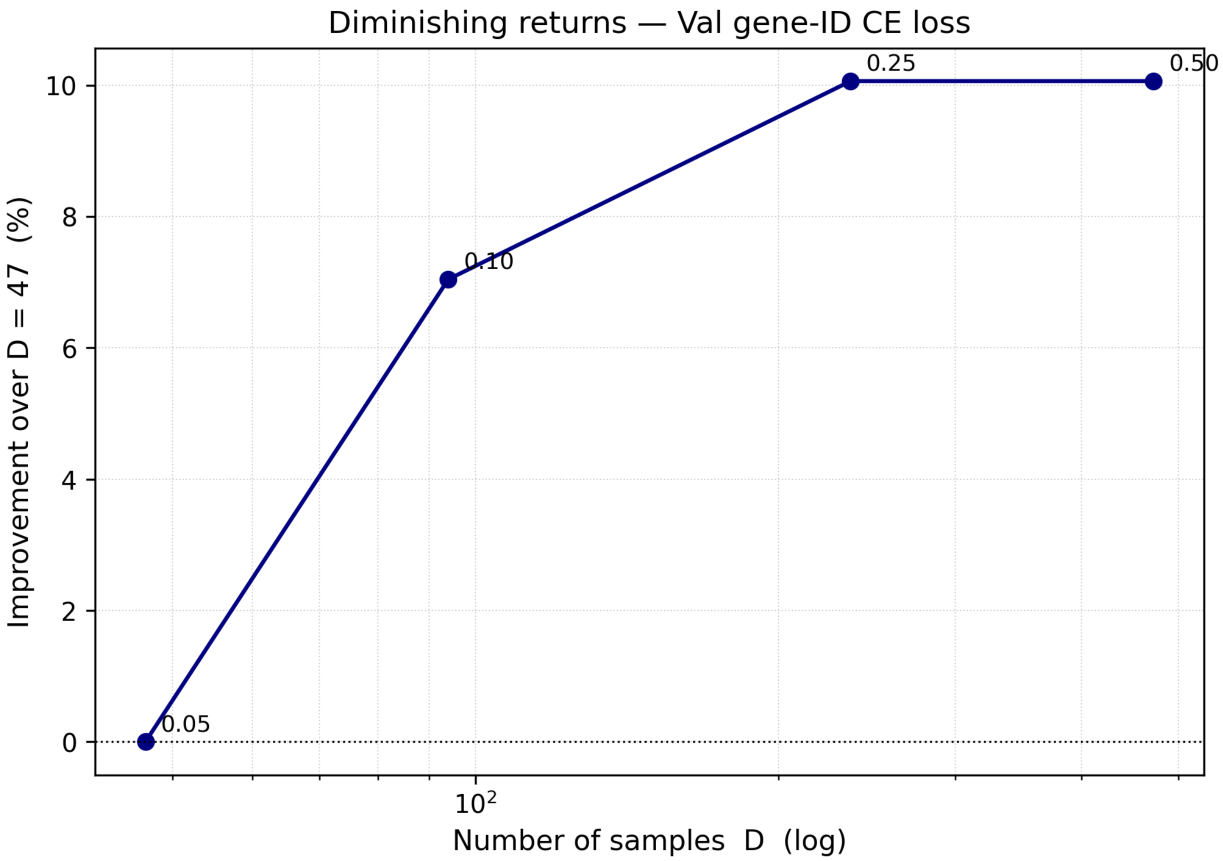

FIGURE S19. **Diminishing returns with increasing pretraining dataset size.** Change in validation gene-ID cross-entropy loss as the number of independent biological samples used for pretraining is increased. The largest improvements are observed at smaller dataset sizes, while increasing the pretraining set from 236 to 472 samples provides little additional improvement.
